# FR-Match: Robust matching of cell type clusters from single cell RNA sequencing data using the Friedman-Rafsky non-parametric test

**DOI:** 10.1101/2020.05.01.073445

**Authors:** Yun Zhang, Brian D. Aevermann, Trygve E. Bakken, Jeremy A. Miller, Rebecca D. Hodge, Ed S. Lein, Richard H. Scheuermann

**Author notes:** **Corresponding Author:** Richard H. Scheuermann.

## Abstract

Single cell/nucleus RNA sequencing (scRNAseq) is emerging as an essential tool to unravel the phenotypic heterogeneity of cells in complex biological systems. While computational methods for scRNAseq cell type clustering have advanced, the ability to integrate datasets to identify common and novel cell types across experiments remains a challenge. Here, we introduce a cluster-to-cluster cell type matching method – FR-Match – that utilizes supervised feature selection for dimensionality reduction and incorporates shared information among cells to determine whether two cell type clusters share the same underlying multivariate gene expression distribution. FR-Match is benchmarked with existing cell-to-cell and cell-to-cluster cell type matching methods using both simulated and real scRNAseq data. FR-Match proved to be a stringent method that produced fewer erroneous matches of distinct cell subtypes and had the unique ability to identify novel cell phenotypes in new datasets. *In silico* validation demonstrated that the proposed workflow is the only self-contained algorithm that was robust to increasing numbers of true negatives (i.e. non-represented cell types). FR-Match was applied to two human brain scRNAseq datasets sampled from cortical layer 1 and full thickness middle temporal gyrus. When mapping cell types identified in specimens isolated from these overlapping human brain regions, FR-Match precisely recapitulated the laminar characteristics of matched cell type clusters, reflecting their distinct neuroanatomical distributions. An R package and Shiny application are provided at https://github.com/JCVenterInstitute/FRmatch for users to interactively explore and match scRNAseq cell type clusters with complementary visualization tools.

## Introduction

Global collaborations, including the Human Cell Atlas [1] and the NIH BRAIN Initiative [2], are making rapid advances in the application of single cell/nucleus RNA sequencing (scRNAseq) to characterize the transcriptional profiles of cells in healthy and diseased tissues as the basis for understanding fundamental cellular processes and for diagnosing, monitoring, and treating human diseases. The standard workflow for processing and analysis of scRNAseq data includes steps for quality control to remove poor quality data based on quality metrics [3–5], sequence alignment to reference genomes/transcriptomes [6–8], and transcript assembly and quantification [8, 9] to produce a gene expression profile (transcriptome) for each individual cell. In most cases, these expression profiles are then clustered [10–13] to group together cells with similar gene expression phenotypes, representing either discrete cell types or distinct cell states. Once these cell phenotype clusters are defined, it is also useful to identify sensitive and specific marker genes for each cell phenotype cluster that could be used as targets for quantitative PCR, probes for *in situ* hybridization assays, and other purposes (e.g. semantic cell type representation where biomarkers can be used for defining cell types based on their necessary and sufficient characteristics [14, 15]).

A major challenge emerging from the broad application of these scRNAseq technologies is the ability to compare transcriptional profiles across studies. In some cases, basic normalization [16, 17] or batch correction [18, 19] methods have been used to combine multiple scRNAseq datasets with limited success. Recently, several computational methods have been developed to address this challenge more comprehensively [20–25]. General steps in these methods include feature selection/dimensionality reduction and quantitative learning for matching. Scmap [20] is a method that performs cell-to-cell (scmapCell) and cell-to-cluster (scmapCluster) matchings. The feature selection step is unsupervised and based on a combination of expression levels and dropout rates, pooling genes from all clusters in the reference dataset. Matching is based on agreement of nearest neighbor searching using multiple similarity measures. Seurat (Version 3) [21, 22] provides a cell-to-cell matching method within its suite of scRNAseq analysis tools. Feature selection is unsupervised and selects highly variable features in the reference dataset to define the high-dimensional space. Both query and reference cells are aligned in a search space projected by PCA-based dimensionality reduction and canonical correlation analysis, to transfer cluster labels through “anchors”. Among many others [23–25], these methods have focused on individual cell level strategies when comparing a query dataset to a reference dataset, not relying on clustering results to guide supervised feature selection or cluster-level matching.

Here, we present a supervised cell phenotype matching strategy, called FR-Match, for cluster-to-cluster cell transcriptome integration across scRNAseq experiments. Utilizing *a priori* learned cluster labels and computationally-or experimentally-derived marker genes, FR-Match uses the Friedman-Rafsky statistical test [26, 27] (FR test) to learn the multivariate distributional concordance between query and reference data clusters in a graphical model. In this manuscript, we first illustrate the matching properties of FR test in this scRNAseq adaptation using thorough simulation and validation studies in comparison with other popular matching methods. We then use FR-Match to match brain cell types defined in the full thickness of human middle temporal gyrus (MTG) neocortex with cell types defined in a Layer 1 dissection of MTG using public datasets from the Cell Types Database of the Allen Brain Map (www.brain-map.org). We also report the cell types that are consistently matched between the two brain regions using multiple matching methods. An R-based implementation, user guide, and Shiny application for FR-Match are available in the open-source GitHub repository: https://github.com/JCVenterInstitute/FRmatch.

## Results

### FR-Match: cluster-to-cluster mapping of cell type clusters

FR-Match, is a novel application of the Friedman-Rafsky test [26, 27], a non-parametric statistical test for multivariate data comparison, tailored for single cell clustering results. FR-Match takes clustered gene expression matrices from query and reference experiments and returns the FR statistic with p-value as evidence that the query and reference cell clusters being compared are matched or not, i.e. they share a common gene expression phenotype. The general steps of FR-Match (Figure 1a) include: i) select informative marker genes using, for example, the NS-Forest marker gene selection algorithm [14]; ii) construct minimum spanning trees for each pair of query and reference clusters (different colors); iii) remove all edges that connect a node from the query cluster with a node from the reference cluster, and iv) calculate FR statistics and p-values by counting the number of subgraphs remaining in the minimum spanning tree plots. Intuitively, the larger the FR statistic, the stronger the evidence that the cell clusters being compared represent the same cell transcriptional phenotype.

**Figure 1.**
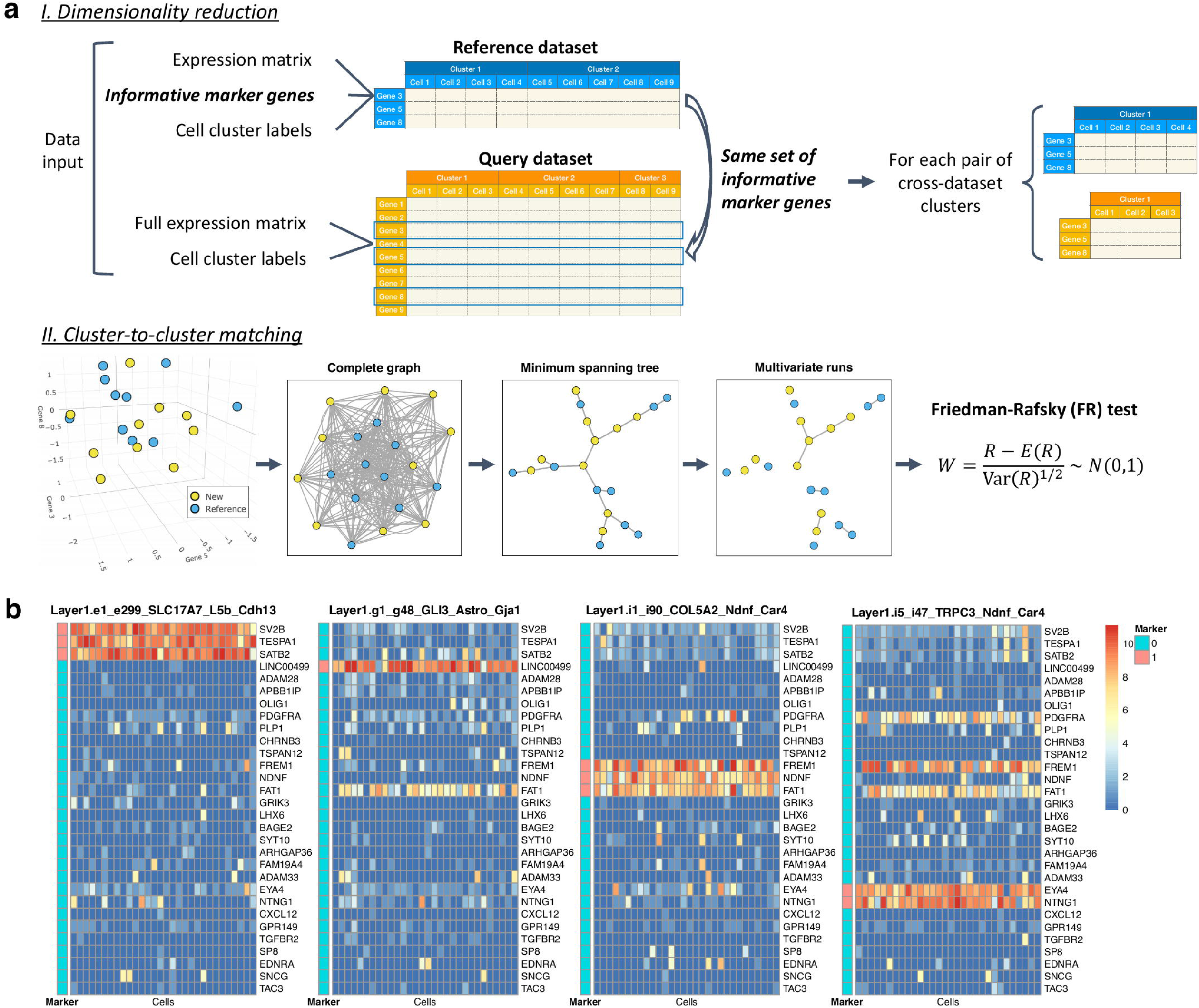
FR-Match schematic and marker gene “barcodes”. **(a)** FR-Match cluster-to-cluster matching schematic diagram. Input data: query/new and reference datasets, each with cell-by-gene expression matrix and cell cluster membership labels. Step I: dimensionality reduction by selecting expression data of reference cell type marker genes from the query dataset. Here, we use the NS-Forest marker genes selected for the reference cell types. Step II: Cluster-to-cluster matching through the Friedman-Rafsky (FR) test. From left to right: multivariate data points of cell transcriptional profiles (colored by cell cluster labels) in a reduced dimensional (reference marker gene expression) space; construct a complete graph by connecting each pair of vertices (i.e. cells); find the minimum spanning tree that connects all vertices with minimal summed edge lengths; remove the edges that connect vertices from different clusters; count the number of disjoint subgraphs, termed “multivariate runs” and denoted as R; calculate the FR statistic W, which has asymptotically a standard normal distribution. **(b)** “Barcodes” of the cortical Layer 1 NS-Forest marker genes in four Layer 1 clusters. Heatmaps show marker gene expression levels of 30 randomly selected cells in each cell cluster. The “Marker” column indicates if the gene is a marker gene of the cluster or not (1=yes, 0=no).

### Supervised marker gene selection provides unique cell type clusters “barcodes”

We adopted the NS-Forest algorithm [14] v2.0 (https://github.com/JCVenterInstitute/NSForest) to select informative marker genes for a given cell type cluster. Applying NS-Forest feature selection to the cortical Layer 1 and full thickness MTG datasets produced a collection of 34 and 157 marker genes that, in combination, can distinguish the 16 cortical Layer 1 [28] and 75 full MTG [29] cell type clusters, respectively. These markers include well known neuronal marker genes like *SATB2*, *LHX6*, *VIP*, *NDNF*, *NTNG1*, etc. (Supplementary Figure 1). The selected marker genes display on-off binary expression patterns producing, in combination, a unique gene expression “barcode” for each cell cluster (Figure 1b). In addition to producing marker genes for each of the individual cell type clusters, this composite barcode serves as an effective dimensionality reduction strategy that captures gene features that are informative for every cell type cluster. The collection of informative marker genes effectively creates an essential subspace that reflects the composite cell cluster phenotype structure in the single cell gene expression data. Thus, supervised feature selection by NS-Forest was used as the dimensionality reduction step for the FR-Match method in this study. Although NS-Forest was used for marker gene selection here, FR-Match is compatible with any feature selection/dimensionality reduction approach that selects informative cluster classification features.

### Matching performance in cross-validation and simulation studies

To assess the performance of FR-Match in comparison with other matching methods, we generated cross-validation datasets utilizing the cortical Layer 1 data and its known 15 cell type clusters for validation studies (excluding the smallest cluster in the original studies with too few cells). Matching was performed using six implementations of the three core methods: FR-Match (using NS-Forest genes), FR-Match incorporating p-value adjustment (FR-Match adj.), scmap (scmapCluster) with default gene selection (500 genes based on dropout proportions), scmap with NS-Forest marker genes (scmap+NSF), scmap with extended NS-Forest marker genes (scmap+NSF.ext) (see Methods section), and Seurat with default gene selection (top 2000 highly variable genes). (Seurat with NS-Forest marker genes was not reported since the results were similar to the results obtained using default marker genes.)

#### Cross-validation assessment of 1-to-1 positive matches

In the two-fold cross-validation study, half of the cells serve as the query dataset and the other half as the reference dataset. Exactly one 1-to-1 true positive match should be identified for each cluster. Figure 2a displays the average matching rate over the cross-validation iterations, where true positives are expected to lay along the diagonal. Four implementations, FR-Match, FR-Match adj., scmap+NSF.ext, and Seurat had excellent performance with 0.93~1 true positive rates (TPR) calculated as the grand average of the diagonal entries. Scmap using its default gene selection approach performed sub-optimally, especially for glial cell types. This is likely due to the fact that informative marker genes for these cell types were not selected using the dropout rate-based feature selection criterion (Supplementary Figure 2). However, using NS-Forest marker genes (scmap+NSF) instead of its default genes resulted in a significant improvement in scmap performance, suggesting that supervised feature selection is advantageous for cell type matching in general. FR-Match implementations had median matching accuracies approaching 0.98 and above, while the next tier performers, scmap+NSF.ext and Seurat, had median accuracies around 0.95 (Figure 2b). Sensitivity and specificity metrics further break down the accuracy measure and indicate the balance between the diagonal (true positive, a.k.a. sensitivity) and off-diagonal (true negative, a.k.a. specificity) matching performance. FR-Match after p-value adjustment is the only algorithm that identified all positive matches. Most methods had very high specificities, whereas FR-Match adj. had somewhat lower specificity due to slightly more false positives.

**Figure 2.**
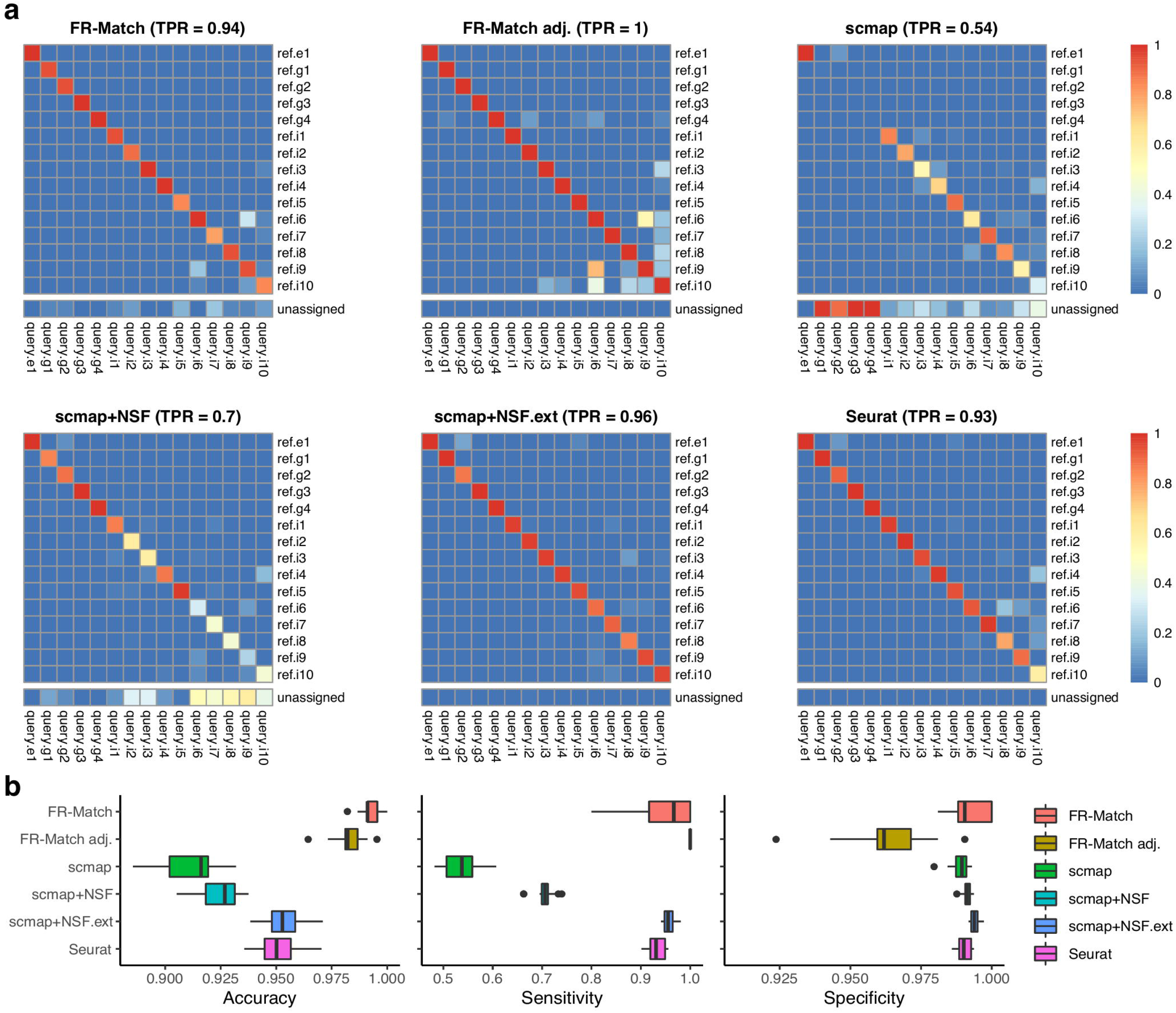
Cross-validation results. Two-fold cross-validation were repeated 20 times on the cortical Layer 1 data with all clusters. Training (reference) and testing (query) data were evenly split in proportion to the cluster sizes. Cluster-level matching results for the cell-level matching methods were summarized as the most mapped cluster labels beyond a defined threshold (see Methods section). Matching output: 1 if a match; 0 otherwise. If a query cluster is not matched to any reference cluster, then it is unassigned. **(a)** Heatmaps show the average matching result for each matching method. True positive rate (TPR) is calculated as the average of the diagonal matching rates, i.e. true positives. **(b)** Median, interquartile range, and full range of accuracy, sensitivity, and specificity of all cluster-matching results in cross-validation for each matching method is shown.

#### Cross-validation assessment of 1-to-0 negative matches

Leave-*K*-cluster-out cross-validation was used to test the performance of these methods under circumstances where one or more cell phenotypes is missing from the reference datasets, i.e. a situation where a novel cell type has been discovered. The left-out cluster(s) should have 1-to-0 match(s) and should be unassigned. While FR-Match implementations clearly identified the left-out cluster as unassigned, other methods produced inappropriate matching when query cell types were missing from the reference dataset (Figure 3). Figure 3a shows results for when the i5 cluster was left out; Supplementary Figures 3-8 show results for when other cluster were left-out in turn. Both FR-Match implementations easily identified the true negative match and correctly labeled the query i5 cluster as unassigned. Other methods partially or primarily mis-matched the query cluster (i5) to a similar yet distinct cluster (i1), as seen in the UMAP embedding where the query i5 nuclei are nearest neighbors to the reference i1 nuclei (Supplementary Figure 9). The accuracy measure for leave-1-cluster-out cross-validation again suggests that the FR-Match method is the best performer with median accuracies approaching 0.99 (Figure 3b). Furthermore, as we removed more and more reference clusters, the FR-Match method showed robust precision-recall curve that consistently outperformed default implementations of scmap and Seurat in ROC analysis (Figure 3c). Seurat’s curve deteriorated because its current implementation lacks an option for unassigned matches; therefore, all cells in the query dataset were forced to map somewhere in the reference dataset. Interestingly, scmap implementations with NS-Forest selected features also had robust precision-recall curves with respect to the increasing number of true negatives.

**Figure 3.**
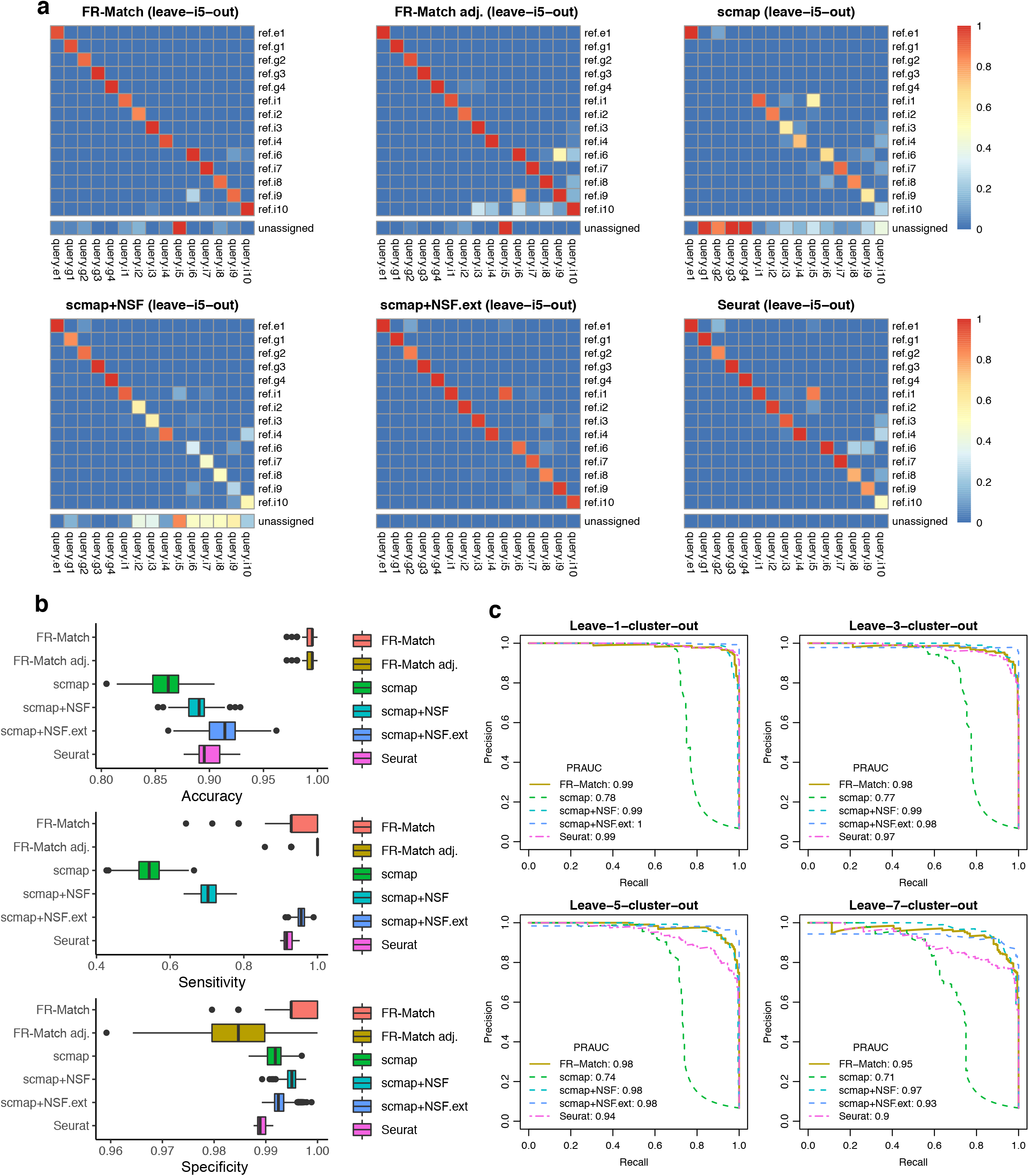
Leave-*K*-cluster-out cross-validation results. The same cross-validation settings as in Figure 2 were used. After data split, *K* ≥ 1 reference clusters were held-out to simulate the situation in which the query dataset contains one or more novel cell type clusters. **(a)** Heatmaps show the average matching result for each matching method when the i5 “rosehip” cluster was left out. **(b)** Accuracy, sensitivity, and specificity of the leave-1-cluster-out cross-validation performance for each matching method is shown. Each cluster was left out in turn, and performance was evaluated across all turns. **(c)** Precision-Recall Curves of the leave-*K*-cluster-out cross-validation performance for *K* = 1, 3, 5, and 7 are shown and Area-Under-the-Curves (AUC) statistics are calculated. Performance was evaluated across 20 iterations of randomly selected *K* clusters. Curves for the FR-Match with and without p-value adjustment have the same shape since the adjustment preserves the order of p-values. Note that the Seurat package by default does not provide for unassigned cells/clusters as a direct output.

The leave-*K*-cluster-out cross-validation has important implications for the capability of each matching method to detect novel cell types in new data sets that are not present in the reference datasets when integrating single cell experiments. In this important use case, the FR-Match method exhibits desirable properties for novel cell phenotype discovery.

#### Simulation of under- and over-partitioning during upstream clustering

Accurate cell type determination from scRNAseq analysis is dependent on accurate partitioning of the cellular transcriptomes into clusters based on their similarity. Existing neuroscientific knowledge [28] suggests that the 15 cortical Layer 1 cell clusters are the current “optimal” clustering of the human brain upper cortical layer scRNAseq data. By combining and splitting these optimal cell type clusters, we simulated under- and over-partitioning scenarios of the upstream clustering analysis. Figure 4a summarizes five cluster partitions ranging from 3 to 18 clusters with F-measure scores indicating the classification power of partition-specific marker genes. The “Top nodes” under-partitioning combines clusters into the three top-level broad cell type classes: inhibitory neurons, excitatory neurons, and non-neuronal cells, producing well known GABAergic, glutamatergic, and neuroglia markers with high F-measure score. The “Mid nodes” under-partitioning combines three groups of closely related GABAergic clusters – i1 + i5, i3 + i4, and i6 + i8 + i9 – resulting in 11 clusters. Over-partitioning of either one (e1) or three (i1, i2, and i3) clusters was performed by running k-means clustering with k = 2 independently for each cluster to simulate real over-partitioning scenarios.

**Figure 4.**
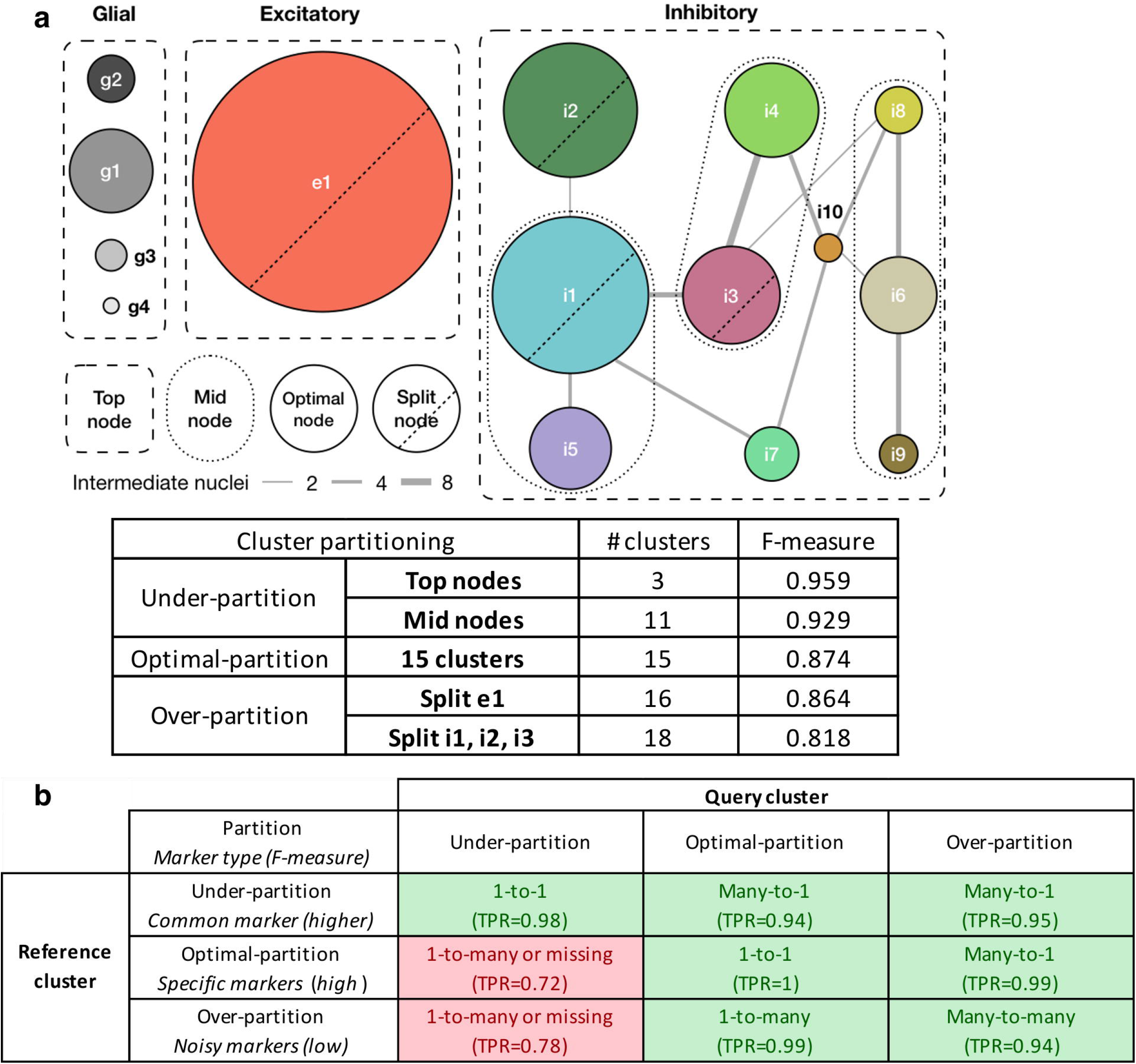
Design of the under-, optimally-, and over-partitioned cluster simulations and their matching properties. **(a)** A schematic of simulating cluster partitions. The optimal partitioning produced nodes where cells were consistently co-clustered across 100 bootstrap iterations for clustering and curated by domain expert knowledge [13, 28]. Connectivity (edge width) between nodes are measured by the number of intermediate cells/nuclei shared by similar nodes. Two under-partition scenarios, “Mid nodes” and “Top nodes”, were simulated by merging similar/hierarchically-connected nodes (e.g. i1 + i5 clusters and all inhibitory clusters, respectively). Two over-partition scenarios, split e1 and split i1, i2, and i3, were simulated by splitting those large size clusters by k-means clustering with k = 2. Median F-measure of the NS-Forest marker genes for each partition are reported in the table. **(b)** FR-Match properties and expected marker gene types with respect to under-, optimally-, and over-partitioned reference and query cluster scenarios, summarized from the simulation results (Supplementary Figure 11). Green blocks in the table are cases with high true positive rate (TPR); red blocks are warning cases with low TPR.

It is important to note that over- and under-partitioning will also have an effect on the gene selection step; it would be predicted that marker gene selection algorithms would have difficulty finding maker genes specific for over-partitioned clusters, which would be reflected in the drop in F-measure scores. Indeed, particularly low F-measure scores may be a good indication of cluster over-partitioning. Figure 4b describes the expected effects on marker gene identification and FR-Match performance after p-value adjustment when clusters are under-, optimally-, and over-partitioned. The types of marker genes that would be selected with different reference cluster partitioning scenarios would impact their ability to effectively drive cluster matching.

Supplementary Figures 10-15 show the matching results of all considered matching methods in various partitioning scenarios. The FR-Match and Seurat methods showed good quality and expected matching results in most partitioning scenarios; scmap had the same problem with the unmatched glial clusters. Seurat showed excellent performance when reference clusters were under-partitioned, but poor performance when query clusters were under-partitioned. Overall, the FR-Match method had stable matching performance in the cluster partitioning simulations. Indeed, 1-to-many and many-to-1 matching results using FR-Match could possibly indicate under-or over-partitioning of the upstream clustering step in scRNAseq data analysis.

#### Simulation of scenarios in which imperfect marker genes are included

Though we recommend using the NS-Forest algorithm to select the minimum set of informative marker genes, users may also want to use their own feature list as the input to FR-Match. There may be other cases where non-informative marker genes have been included. In order to assess the performance of FR-Match with respect to less than ideal marker gene lists, we use simulation to evaluate the matching performance in two scenarios: i) when there are non-informative (i.e. noisy) genes in the features selected, and ii) when some informative marker genes are missing from the feature list with or without non-informative genes. Throughout this simulation study, the FR-Match adj. implementation was used.

To simulate scenario (i), we used the 32 NS-Forest marker genes associated with the 15 cell types in the Layer 1 data, together with randomly selected genes from the 16,497 available genes in the dataset. In this scenario, the barcoding pattern of the informative marker genes were preserved, whereas the random genes showed more noisy and non-specific expression patterns in the “barcode” plots (Supplementary Figure 16a). In the simulations, we increased the number of extra genes added from 1 to 15; FR-Match was very robust to noisy genes in each simulated case with true positive rate staying close to 1 (Supplementary Figure 16b). Other performance measures – accuracy, sensitivity (true positive rate), and specificity (true negative rate) – all stayed well-above 0.9, suggesting that the overall performance of FR-Match was stable and robust, even when the marker gene list contained up to 30% non-informative genes (15 extra genes) (Supplementary Figure 16c). Increasing the number of non-informative genes may slightly impact the specificity due to more false positives (off-diagonal intensities in Supplementary Figure 16b) and therefore leads to the slight downward trend of the overall accuracy.

For simulation scenario (ii), we generated two subcases to illustrate the impact of interfering with different combinations of marker genes on the matching performance. In the first subcase, we removed marker genes for three very distinct cell types: an excitatory cell type (e1), a glial cell type (g1), and an inhibitory cell type (i1); and used the remaining NS-Forest marker genes to match all cell types in the Layer 1 dataset. Surprisingly, each cell type was matched correctly most of the time with an overall true positive rate of 0.98 (Supplementary Figure 17a). We also replaced the removed marker genes with the same number of random genes; the matching performance was also very good, and the impact of the changes in the marker gene list was insignificant (Supplementary Figure 17a). In the second subcase, we considered removing/replacing the marker genes for two related inhibitory cell types: i1 and i2. Without marker genes that distinguish these similar cell types, FR-Match matched the i1 and i2 cell types to each other (i.e. a many-to-many match) while maintaining the distinction from other cell types with informative classification markers (Supplementary Figure 17b). The “barcode” plots for i1 and i2 became generally non-selective with random expression of some other inhibitory markers in the background (Supplementary Figure 17c). Such indistinct “barcode” plots may be an effecting warning for many-to-many matches. The absence of good classification markers is most harmful to specificity (due to false positives), while sensitivity (true positive rate) remains high (Supplementary Figure 17d).

In summary, as long as informative marker genes with good classification power are selected, FR-Match is robust to other non-informative genes included in the feature list. Many-to-many matching results by FR-Match may be a good indicator of the absence of informative marker genes between the mis-matched cell types.

### Cell type mapping between cortical Layer 1 and full MTG

We next extended the validation testing to a more realistic real-world scenario where a new dataset has been generated in the same tissue region using slightly different experimental and computational platforms. We tested FR-Match with p-value adjustment using two single nucleus RNA sequencing datasets from overlapping human brain regions – the single apical layer of the MTG cerebral cortex (cortical Layer 1), in which 16 discrete cell types were identified [28], and the full laminar depth of the MTG cerebral cortex, in which 75 distinct cell types were identified [29]. We selected NS-Forest combinatorial marker genes separately for each dataset. The marker gene sets may contain overlapping genes for some cell types, e.g. *CUX2* is a useful marker gene for more than one layer 2-3 cell types in combination with other marker genes; classification power of these combinatorial marker genes are evaluated in detail in another study [30].

Matching results were assessed from two perspectives: i) agreement with prior knowledge such as layer metadata from the design of these experiments [28, 29], and ii) agreement with other matching methods. Since these datasets targeted the same cortical region with overlapping laminar sampling, we expect that matching algorithm should find 1-to-1 matches of each cell types in the cortical Layer 1 data to one cell type in layers 1-2 from the full MTG data. The final matching results were concluded from two matching directions: Layer 1 query to MTG reference with MTG markers, and MTG query to Layer 1 reference with Layer 1 markers. The two-way matching approach was applied to all comparable matching algorithms.

#### FR-Match uniquely maps cell types reflecting the overlapping anatomic regions

Using FR-Match, we mapped each of the 13 Layer 1 cell types uniquely to one MTG cell type (Figure 5a), i.e. 1-to-1 two-way matches. These matches precisely reflect the overlapping anatomic regions in these two independent experiments in that the matched MTG cell types all have an “L1” layer indicator in their nomenclature. The one exception for the Layer 1 e1 cluster likely reflects the incidental capture of upper cortical layer 2 excitatory neurons in the original Layer 1 experiment [28]. And while most of the *SST* cell subtypes are located in deeper cortical layers, FR-Match specifically selected the small number of L1 *SST* clusters as top matches. The same was true for *VIP* and *LAMP5* cell subtypes. The minimum spanning tree plots produced by FR-Match provide a clear visualization of matched and unmatched cell clusters (Figure 5b).

**Figure 5.**
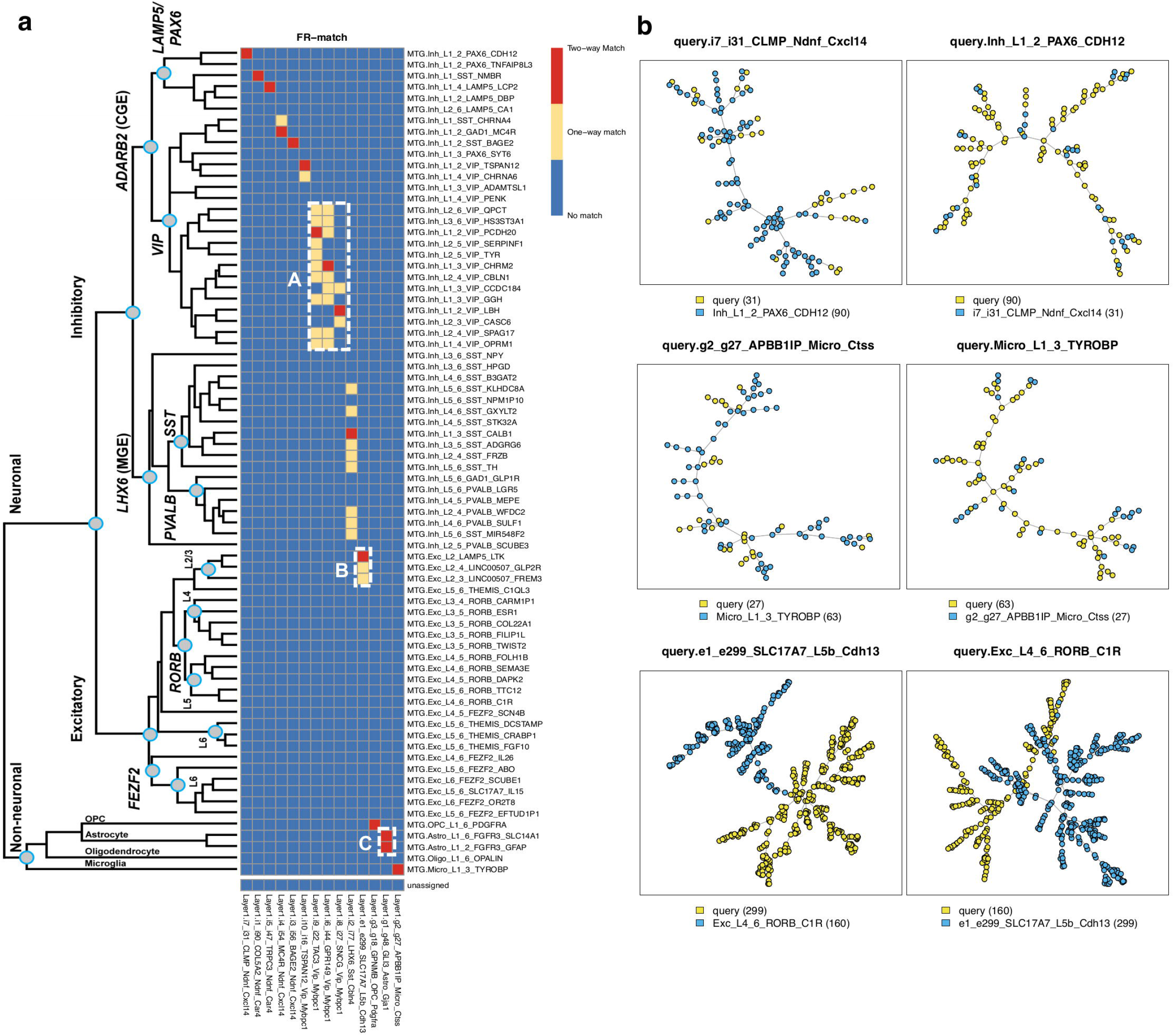
FR-Match results for cell type matching between the cortical Layer 1 and full MTG datasets. **(a)** Two-way matching results are shown in three colors: red indicates that a pair of clusters are matched in both directions (Layer 1 query to MTG reference with MTG markers, and MTG query to Layer 1 reference with Layer 1 markers); yellow indicates that a pair of clusters are matched in only one direction; and blue indicates that a pair of clusters are not matched. The hierarchical taxonomy of the full MTG clusters is from the original study [29]. FR-Match produced 13 unique, and two non-unique two-way matches between the two datasets. Box A shows densely located one-way matches in the subclade of *VIP*-expressing clusters. Box B shows incidentally captured cells from upper cortical Layer 2 mixed in the Layer 1 e1 cluster. Box C shows the non-unique two-way matches of astrocyte clusters. **(b)** Examples of matched and unmatched minimum spanning tree plots from the FR-Match graphical tool. Top row: examples of two-way matched inhibitory clusters. Middle row: examples of two-way matched non-neuronal clusters. Bottom row: examples of unmatched excitatory clusters from different layers. Legend: cluster name (cluster size).

To validate further, we compared the matching results to the hierarchical taxonomy of MTG cell types [29], which reflects cell type relatedness (left side of Figure 5a). First, the block of one-way matches in Box A precisely corresponds to a specific sub-clade of *VIP*-expressing cells with close lineage relationships, suggesting that one-way FR-Match results are evidence of closely related cell types. Second, FR-Match correctly identified excitatory neurons that were incidentally captured from upper Layer 2 in the cortical Layer 1 experiment in Box B, corresponding to L2/3 excitatory neurons in the full MTG dataset. Third, Box C suggests under-partitioning of the Layer 1 astrocyte cluster as multiple two-way matches were found for the same cluster.

Directional one-way matching results are shown in Supplementary Figure 18. Though different matching patterns are observed from each direction, they reflect the fact that these datasets are measuring different cell types. There are some cases where the difference might be due to the cell complexity in the datasets, e.g. the *VIP* or *SST* types, and this might be leading to the dynamic range and skewness of p-value distributions for each query cluster.

#### Cell type mapping using other existing approaches

In mapping cell types between cortical Layer 1 and the full MTG, both FR-Match and Seurat produced similar unique two-way matches (Figure 6). Examining all matching results and all matching algorithms, FR-Match produced the most “conservative” mapping of cell types. The other matching algorithms produced matching results that had more sparsely-distributed *VIP* types (Box A), and were not laminar specific (Box B). Among all approaches, glial cell types were mapped somewhat differently (Box C), probably due to their overall lower sampling and distinct phenotypes compared to the majority of GABAergic and glutamatergic neurons.

**Figure 6.**
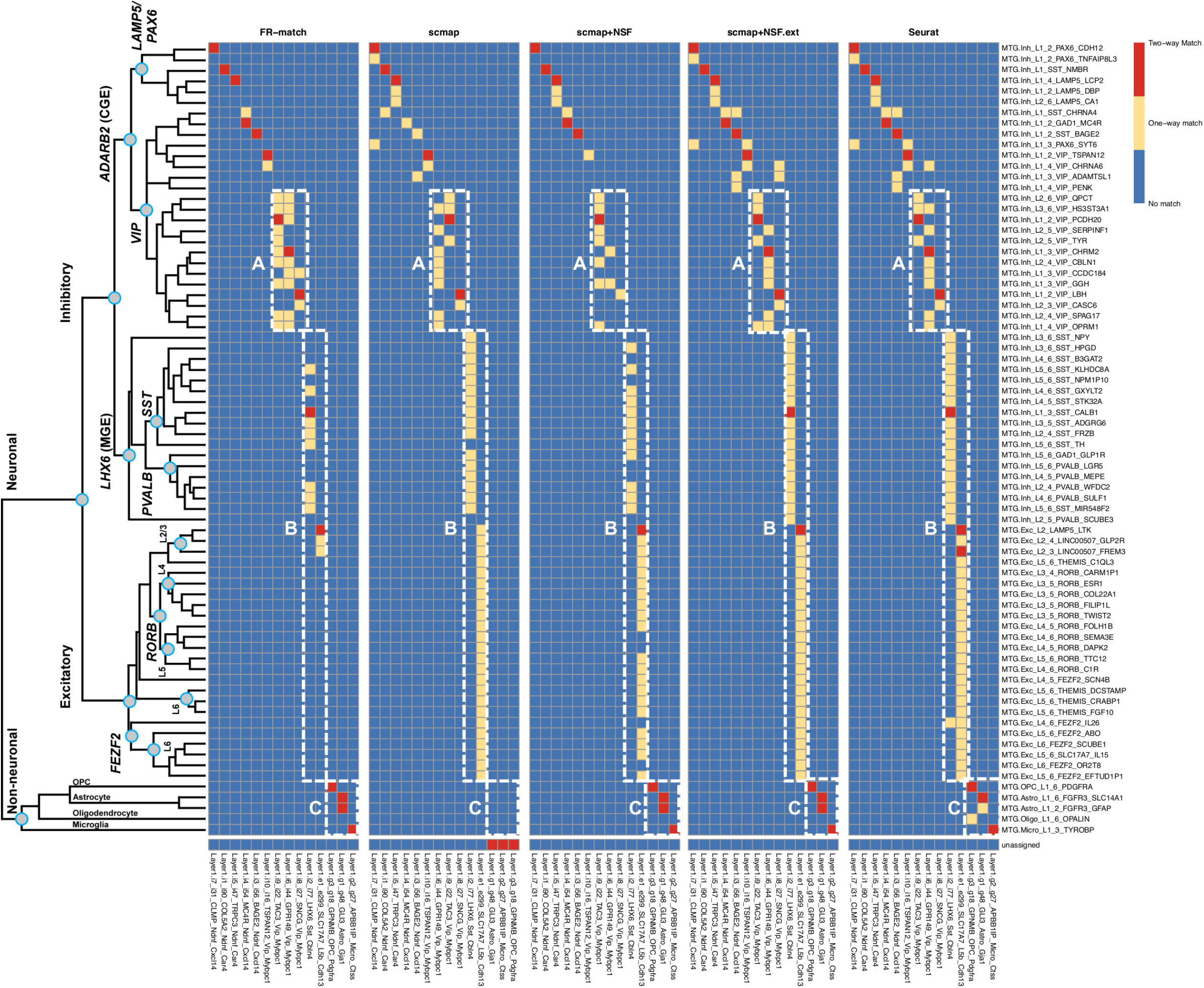
Cell type matching results between the cortical Layer 1 and full MTG datasets using other matching methods. Two-way cluster-level matching results for the cell-level matching methods were summarized as the most mapped cluster labels beyond a defined threshold (see Methods section). Box A shows matches in the *VIP*-expressing subclade. Box B shows matches spanning multiple layers among the MTG clusters. Box C shows matches of glial clusters.

FR-Match shows three advantages over the alternative methods. First, by using supervised feature selection for each cell type, major and minor cell populations are equally represented in the reduced-dimensional space for cell type matching. This strategy would also benefit other matching methods with sub-optimal feature selection/dimensionality reduction. Second, FR-Match clearly excludes the matching of cell types that are only present in one of the datasets. Third, FR-Match allows one-to-multiple and unassigned matches, which allows for detecting potential cluster partitioning issues and the discovery of novel cell types.

The other existing cell-level matching approaches naturally provide the probabilistic cluster-level matching of cell types as the percentage of matched cells in query cluster (Supplementary Figures 19-22); a deterministic cluster-level match would depend on the selection of an *ad-hoc* cutoff of the probabilistic matching. Thus, deterministic cell type mapping or discovery of novel cell types would be difficult as i) individual cells may be alike in the same broad cell class even if the specific cell type may not be present in the reference dataset, and ii) the probabilistic cutoff may be subjective. Therefore, both scmap and Seurat identified many more non-specific one-way matches than FR-Match, which uses an objective p-value cutoff. Combining all results, we finally report 15 high-confidence ensemble matches between Layer 1 and full MTG cell types in Supplementary Table 1.

#### The effects of alternative gene selection and cell clustering methods on matching performance

To further elucidate the impact of alternative gene selection or cell clustering choices on cluster matching, we performed the following analyses.

In the two brain datasets, cell types are defined and characterized by a domain knowledge-guided iterative clustering [13] and transcriptomically-derived markers [28, 29]. The nomenclature used to describe these cell types consists of the broad cell class (inhibitory, excitatory, and glial cells), layering information (for the MTG dataset), one marker gene for the subclass node in the taxonomy tree (e.g. *VIP*, *SST*, etc.), and one marker gene for the leaf node cluster. For example, the “Inh_L1_2_PAX6_CDH12” from the MTG dataset means the inhibitory neurons located in layer 1-2 within the *PAX6*-subclass/subbranch expressing *CDH12*. The leaf node marker genes are preferentially selected by a binary scoring scheme [29] different from the one used by NS-Forest. Thus, the “cell type naming genes” provide an alternative informative marker gene set.

To assess matching performance using a different set of informative marker genes, we replaced the NS-Forest marker genes by these cell type naming genes for both datasets, followed by the same matching approaches. 26 and 87 naming genes were defined for the Layer 1 and full MTG datasets, respectively, out of which, 9 and 18 genes are in common between the naming genes and the NS-Forest marker genes, respectively. Using cell type naming genes, FR-Match, scmap, and Seurat all performed slightly differently with less ideal matching patterns (Supplementary Figure 23). Overall fewer matches were identified; and the identified matches were less specific (i.e. mapping to neighboring cell types). This is probably because using only one leaf node marker gene may not be enough to fully capture the differences between those closely related leaf node cell types. From these matching results, we may conclude that NS-Forest selects better sets of informative markers than the other approach in this example, which has an impact on all three matching methods; less optimal feature selection will negatively impact matching regardless of the matching methods.

In another analysis, we compared the matching performance of FR-Match, scmap, and Seurat with respect to a different clustering method. The community detection Louvain method [10] is one of the most commonly used clustering methods for scRNAseq analysis. We applied Louvain clustering (implemented in the Seurat R package, with resolution = 1) to the full MTG dataset, which resulted in 26 reasonably segregated clusters in the UMAP low-dimensional embedding space (Supplementary Figure 24a). Matching results with the Louvain clusters are shown in Supplementary Figure 24b. FR-Match produced similar matching results regardless of the clustering methods: each Layer 1 cluster is strongly matched (two-way match) to some Louvain cluster of the full MTG dataset. Many-to-one and one-to-many matches are observed since the generic Louvain method appears to have under-partitioned the data in comparison with the original expert-curated iterative clusters, which agrees with the matching patterns we observed in our simulations. Matching by scmap and Seurat with the Louvain clusters shows the same problems as with the original clusters, i.e. excessive unassigned matches (scmap), and non-specific matches of the Layer 1 excitatory cluster (scmap and Seurat). Using different clustering methods will lead to different matching results depending on the clustering quality. As long as the clusters are reasonably good, FR-Match is able to detect high quality matches regardless of the clustering methods.

#### Cell type matching using batch integration

To date, there are more than 10 methods that have been proposed to correct the batch effects of scRNAseq data; among them, Harmony [31], LIGER [32], and Seurat 3 [21] are the recommended algorithms for batch integration [33]. Only Seurat is an end-to-end pipeline that inputs multiple scRNAseq data batches and outputs cell-to-cell alignment between batches. By summarizing the cell-level batch integration with prior cluster memberships of the cells, we compared the performance of Seurat for cell type matching with FR-Match in previous subsections. In this subsection, we implemented a workaround for Harmony and LIGER to transfer the batch integration outputs to produce putative cell type matches.

We applied Harmony (Supplementary Figure 25-26) and LIGER (Supplementary Figure 27-28) individually to integrate the Layer 1 and MTG datasets; both methods showed effective “batch-effect” removal in the UMAP (Supplementary Figure 25b-c) or tSNE (Supplementary Figure 27b-c) low-dimensional embedding. For both Harmony and LIGER, the outputs from the algorithms are the integrated cells in some dimensionally reduced spaces; joint clustering can then be conducted on the integrated data spaces (Supplementary Figure 25d, Supplementary Figure 27d); and cell type matching can be inferred from the “river” plots (Supplementary Figure 26a, Supplementary Figure 28a) between the input batches through the common joint clusters. We transferred the river plot to a one-to-one correspondent cell type matching heatmap, with each match indicating there exists a path between the two cell types in the river plot. Note that the heatmap is non-directional for a given set of edges of the river plot. Through such a workaround, we obtained cell type matching results for Harmony (Supplementary Figure 26b) and LIGER (Supplementary Figure 28b) in a similar format as FR-Match. It is clear that the batch integration approaches produce matches in blocks (i.e. many-to-many matches), and do not effectively yield the specific matches within these blocks if multiple related cell subtypes are presented. These batch integration methods were not originally designed for the task of cell type integration; therefore, it is not surprising that they produce sub-optimal results.

## Discussion

FR-Match offers a cluster-level approach for mapping cell phenotypes identified in scRNAseq experiments. It extends the current cell-level matching algorithms by: i) borrowing information from all the cells in the same cluster using a statistical test that provides both probabilistic matching in p-values and objective p-value thresholds for deterministic matching, and ii) providing simple visualization of cell type data clouds in the minimum spanning tree graphical representation. Matching results of FR-Match are relatively conservative yielding highly specific matches, which can confirm cell type equivalence, lead to novel cell type discovery, and diagnose upstream clustering problems. Among many other scRNAseq data integration strategies, this approach combines informative feature selection and cluster-level integration of the NS-Forest and FR-Match software suites, producing intuitive results with high interpretability, including useful intermediate results such as binary marker genes and minimum spanning tree graphs for users to monitor and gain meaningful insights from the mapping solutions.

Based on the computational and statistical investigation of both simulated and real datasets, we conclude that: i) the FR-Match and Seurat methods show excellent performance in mapping neuronal and glial cell types using snRNAseq data from human brain; and ii) supervised feature selection, such as the NS-Forest algorithm, appears to produce excellent marker gene combinations that can be used as an effective feature selection/dimensionality reduction technique for cell type mapping with multiple methods, including FR-Match and scmap. Scmap is a consensus method that requires at least two of the three association metrics – cosine similarity, Pearson and Spearman correlations – to be in agreement as the last step to determine a match, thus the comparative analysis results of the matching methods reported here may also serve as a reference guide for matching performance using those association metrics.

One of the biggest challenges in scRNAseq alignment at the moment seems to be the proper assignment of cells from a cell type found in only one dataset. These cells are often matched to a closely related cell type in a second dataset. In this regard, FR-Match appears to be superior in being able to determine which cell types from two datasets are *not* matched, an indication of novel cell type discovery.

For all compared methods in this study, it’s interesting to note that under-partitioning the query clusters leads to degraded performance, unless the reference clusters are also under-partitioned. This suggests that a useful strategy would be to map to reference types in a hierarchical manner by first mapping to broad classes of references types and then moving down the tree to finer types until ambiguous matches appear. The negative effect of under-partitioned clusters also applies to the nested classes of heterogeneous cell types.

Currently, decisions on splitting a cell type class or joining two subclasses are often determined by subjective examination of the expression distributions of selected gene expression markers. In this regard, one use of FR-Match could be to provide for an objective assessment of cell clusters partitioning when comparing a new query dataset to a high-quality reference dataset from a carefully curated cell atlas knowledgebase. Assuming the reference cell types are optimally partitioned, a 1-to-many match using FR-Match would suggest that the query cluster is under-partitioned and should be further split into sub-clusters; a many-to-1 match would suggest that the query clusters are over-partitioned and should be merged. In this use case, FR-Match can be used as a tool to guide optimal partitioning of scRNAseq data clustering, leveraging information captured in well-curated cell type reference knowledgebases.

Automated cell type integration of independent scRNAseq datasets remains challenging. Creating an unbiased, high-resolution and comprehensive cell type reference would be a critical task for the whole single cell research community. Consensus mapping schemes that survey both cell-level and cluster-level matchings will be useful for establishing such a reference data atlas. We believe that final mapping of the brain cell types agreed upon by the type of bi-directionally and multi-level matchings reported here represents the best-practice for computational cell type mapping, requiring minimal expert intervention.

Single cell evaluation is a fast-evolving field. Although not fully explored here, we expect FR-Match to be applicable to cross-platform, cross-specimen, cross-anatomy, and cross-species matching of scRNAseq clustered data. The effect of dropouts and the dynamic range of single cell sequencing data from protocols other than the Smart-seq [34] protocol stand out as key challenges to be overcome. To address these challenges, we are now developing add-on features to the core FR-Match algorithm, including imputation techniques [35] for the relatively high dropout rates in 10X Genomics droplet-based protocols [36], and moment-based normalization options [37] for the discrete and dispersed values produced in single cell spatial *in-situ* hybridization protocols [38–40]. Preliminary results of mapping Smart-seq cell clusters to 10X cell clusters suggest that FR-Match will be useful for cross-platform cell type matching when appropriate dropout imputation and data normalization upstream steps are included in the computational pipeline (data not shown). While these emerging technologies will produce more complicated data integration challenges, the adaptation of methods like FR-Match are poised to play an essential role in the broad integration of scRNAseq cell phenotyping experiments.

## Methods

### The cell type matching problem

Consider two single cell RNA sequencing experiments – one query/new experiment and one reference experiment. A cell-by-gene expression matrix for each experiment is obtained by standard scRNAseq data processing and analysis workflows, including quality control, reference alignment, sequence assembly, and transcript quantification. Cell cluster labels are also obtained from clustering analysis using, for example, the community detection Louvain algorithm [10], and/or other domain specific knowledge. These cell clusters represent transcriptionally-distinct cellular phenotypes within each experiment. The cell type matching problem is whether a pair of query and reference cell clusters identified in related but independent experiments are instances of the same or different transcriptionally-defined cell phenotypes.

We propose a computational solution to the cell type matching problem – FR-Match – an adaptation the Friedman-Rafsky statistical test for scRNAseq data, which takes two input datasets (query and reference) each with a gene expression matrix and cell cluster membership labels (Figure 1a). Importantly, FR-Match uses a set of informative marker genes that characterize the reference cell type clusters. Dimensionality reduction is done by imposing the same set of marker genes on the query dataset, to select the most informative features shared with the reference dataset. For each pair of cross-dataset clusters, we perform cluster-to-cluster matching via the Friedman-Rafsky statistical test. As a result, FR-Match outputs the following types of match (format: query-to-reference): 1-to-0 or unassigned (indicative of a novel cell type), 1-to-1 (indicative of a uniquely matched cell type), 1-to-many (indicative of an under-partitioned query cluster or over-partitioned reference cluster), many-to-1 (indicative of an over-partitioned query cluster or an under-partitioned reference cluster).

### Necessary and sufficient marker gene identification by random forest

In order to perform dimensionality reduction, random forest machine learning as implemented in the NS-Forest algorithm [14, 15, 30] (v2.0 at https://github.com/JCVenterInstitute/NSForest) was used to select necessary and sufficient marker genes for each reference cell type cluster. NS-Forest includes steps for: i) feature selection, ii) feature ranking, and iii) minimum feature determination. Let *X* be an *n* x *m* dimensional cell-by-gene matrix, where n is the number of cells and *m* is the number of genes. Let y be an *n* x 1 vector of cluster labels. In step (i), random forest models, with 10,000 decision trees each, are built for input data *X* and each cluster label in *y* under a binary classification scheme. From each random forest model, the average information gain based on the Gini index for each gene is extracted, which is then used as a measure of feature importance to rank the gene features. In step (ii), for the top 15 ranked genes, a binary expression score for gene *g* in cluster *k* is calculated as

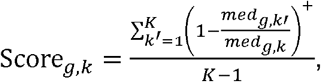

where *med_g,k_* is the median expression level of gene *g* in cluster *k*, *K* is the total number of clusters, and (·)^+^ defines the non-negative value of the equation. The binary expression score ranges from 0 to 1, where 1 indicates absolute binaryness, i.e. the gene exclusively expressed in the target cluster and not at all in non-target clusters. In step (iii), the top 6 genes from step (ii) are selected and all combinations are evaluated by the F-beta score. F-beta is an F-measure weighted by *β* such that

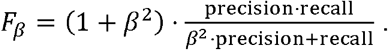

*β* = 0.5 was set to weight precision more than recall, which compensates the effect of false negatives dropouts due to technical artifacts in scRNAseq experiments. The output from step (iii) is a minimum set of marker genes for each cell type cluster (usually 1 – 4), whose expression in combination is sufficient to discriminate the target cell type cluster from the rest of the cells. In addition to the minimum set of NS-Forest marker genes, the algorithm also provides an extended list of binary marker genes as a supplementary output from step (ii), which may achieve higher discriminative power under certain circumstances. The top 15 NS-Forest genes for each cell type formed an NS-Forest extended gene list as an alternative feature selection option for matching algorithms. For a more detailed discussion of the choice of the number of top genes used in NS-Forest v2.0, see Aevermann et al. [30]

### Friedman-Rafsky test

The Friedman-Rafsky (FR) test [26] is a multivariate generalization of the non-parametric two-sample comparison problem. This classical statistical test is distribution free. Consider two general distributions *F_x_* and *F_Y_* for samples (*x*_1_,…, *x_m_*) and (*y*_1_,…, *y_n_*) in a *k*-dimensional space, respectively. (In the context of FR-Match, the *x*’s and *y*’s denote the expression profiles of each cell in the query and reference clusters; *m* and *n* are the number of cells in each cluster; *k* is determined by the number of informative marker genes from the reference dataset). Under the hypothesis testing framework, the original FR test is designed for testing

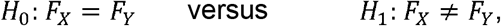

in which the null hypothesis states that the cells from both query and reference clusters are from the same transcriptional distribution; the alternative hypothesis states that the two cell populations are from different transcriptional distributions. Thus, the cell type matching problem becomes a statistical test to detect comparisons for which *H*_0_ is true.

The underlying model of the FR test is a graphical model based on the minimum spanning tree of pooled samples (Figure 1a). In the multi-dimensional informative marker gene space, cells from different clusters (indicated by colors) are pooled and form a mixture of data points. A complete graph can be constructed, which connects all cells to each other and uses the edge length to preserve the pairwise Euclidean distance between cells in the original space. Next, the complete graph is trimmed to a tree graph that connects all cells with the minimum total length of edges, i.e. the minimum spanning tree. Edges that connect cells of different clusters are then removed and the number of disjoint subtrees is counted. Intuitively, if there are a large number of subtrees, it implies that the pooled cells are closely interspersed and therefore more likely to be from the same multivariate gene expression distribution.

Formally, let *R* be the total number of subtrees – “multivariate runs” in the FR test framework, with mean *E*(*R*) and variance *Var*(*R*) directly derived from graph theory. The FR statistic is defined as

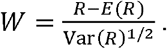

Friedman and Rafsky showed that the asymptotic distribution of W follows a standard normal distribution for large sample sizes:

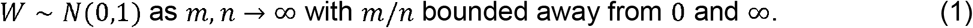

For the hypothesis testing purpose, *H*_0_ is rejected for small values of *W*, i.e. p-value is one-sided such that *p* = Pr(*W* ≤ *w*). Note that, in the cell type matching application, we determine a *match* if *p* > 0.05, but other p-value thresholds could also be used.

### FR-Match method

Extending from the classical statistical test, FR-Match is a novel application of FR test to approach the cell type matching problem with scRNAseq data. The full FR-Match algorithm not only implements the basic testing procedure, but also adapts modifications for specific issues pertaining to the scRNAseq application. A major issue is that two cell clusters to be compared may have very different cluster sizes, such as a dozen cells versus hundreds of cells (Supplementary Figure 29). The unbalanced cluster sizes will often cause two problems: i) unstable statistical power as the ratio of cluster sizes deviates from the asymptotic condition, and ii) exponentially long computational time needed for constructing minimum spanning tree for large number of cells. To address these problems, an iterative subsampling scheme was implemented, which repeatedly performs sampling without replacement of *S* cells from each cell cluster for *B* times. (For clusters with fewer than *S* cells, sample all cells in the cluster.) Default values of *S* and *B* are 10 and 1000, respectively, but are tunable. The median p-value of all iterations is outputted. Other modifications include filtering small clusters with less than C cells each, and p-value adjustment for multiple hypothesis testing correction. Empirically, *C* = 10 was chosen for defining a cell type cluster with high confidence since it appeared to provide enough cell instances to be representative. It is suggested to set *S* = *C*, but it is not a necessary condition for the algorithm. A disproportionate ratio of *S* to *C* would adversely affect the underlying statistical assumptions due to the unmet asymptotic condition in Equation (1).

As an alternative to the asymptotic theory, permutation testing is a widely-accepted practical choice for approximating the null distribution of the FR statistic in a hypothesis testing framework [41]. We designed a simple technical simulation to compare the statistical properties of the FR test, FR permutation test, and FR test with subsampling scheme, with respect to the major pragmatic concern of imbalanced cluster sizes that specifically pertains to the cell type matching problem. We generated multivariate data from a Multivariate Normal (MVN) distribution (k = 40 dimensions). Random samples were drawn from (*x*_1_,…, *x_m_*) ~ *MVN*(*μ* = 0, Σ = *I*) and (*y*_1_,…, *y_n_*) ~ *MVN*(*μ* = 0 + δ, Σ = *I*), where is the identity matrix. Under the null, δ = 0, i.e. no location difference between the x- and y-samples; under the alternative, we set δ = 0.4 for moderate location shift in their distributions. To simulate the imbalanced cluster sizes, we fixed one cluster size *m* = 10 and varied the other cluster size *n* = 10, 20, 100, 200. The ROC analysis (Supplementary Figure 30) confirm that the permutation test is a very good approximation of the FR test based on asymptotic theory; however, both tests show deteriorating ROC curves when the sample sizes were very imbalanced (*n* = 200, blue curve). In contrast, FR test with subsampling shows the most ideal property – better ROC curve and larger AUC value – as sample size (i.e. cluster size in this context) increases. Therefore, the iteratively subsampling scheme was adopted in the FR-Match algorithm.

In our proposed procedure, the subsampling parameter *S* was initially chosen based on practical considerations, we also provide more simulation results for guiding the choice of *S* here. Based on the same simulation design as above, we evaluated the AUC values for FR subsampling tests with *S* = 10, 20, 30, and benchmarked with the FR test (Supplementary Figure 31). When both input cluster sizes m and n vary from 10 to 200, the FR subsampling test with *S* = 10 outperforms all other choices with the FR test showing the highest AUC values in all simulated cases with *m* and *n*. This is potentially due to the expectation that the choice of *S* should embrace the right balance between gathering enough samples to represent the whole cluster and avoiding local structures in the cluster (i.e. large subtrees of the same color in an MST). We believe this might be related to the “effective” dimensionality of the data space characterized by Σ and other distributional properties, which will be an interesting topic for future statistical research. In this manuscript, the choice of *S* is supported by empirical evidence; readers should use their own judgement on the choice of *S* for their own datasets. Though fixed-size subsampling may result in increasing variability for larger clusters [42], our adapted procedure of the FR test is a pragmatic solution to the imbalanced cluster size issue. Developing a general solution to solve this unmet statistical assumption problem is beyond the scope of these studies.

In the Layer 1 and full MTG matching analysis reported in this manuscript, tunable parameters were set at the default values described above. When a sequence of FR-Match p-values were computed for each pair of Layer 1 cell type and MTG cell type, Benjamini & Yekutieli [43] p-value adjustment was applied for multiple hypothesis testing correction before the final determination of a cell type match.

### Determining cluster-level match for the cell-level matching methods

In comparison with other popular matching methods, a voting rule was adopted after obtaining the cell-level matching results from algorithms scmap (cell-to-cluster) and Seurat (cell-to-cell). Scmap provides a map: query cell → reference cluster. We calculate the % of reference cluster labels grouped by the query cell labels, and thereby obtain a quantitative measure ranging from 0 to 1 that indicates the probability of being the same cell type between the query and reference cell clusters. Similarly, the Seurat alignment is extended to query cell → reference cell → reference cluster, and calculate the cluster-to-cluster matching measure in the same way. For a specific query cluster, its cluster-level match is determined by the votes of its member cells for their mapped reference cluster labels. An ad-hoc threshold at 30% was used for defining a deterministic match, which accounts for both the detection of a substantial proportion of query cells matched to one reference cluster and the possibility that some query clusters might be matched to multiple reference clusters. If the 30%-criterion is not met, then the query cluster is defined as unassigned in the matching results. The cluster-level matching results may change depending on the ad-hoc threshold used. For example, if changing the threshold to 40%, Seurat would identify the same set of two-way matches, but with three fewer one-way matches (Supplementary Figure 32). A data-driven decision on such a threshold can be guided by the distribution of % of matched cells in Supplementary Figures 19-22.

### Cross-validation and simulation design

Data generation for the cross-validation and simulation studies were from the cortical Layer 1 data with 15 cell clusters [28] (excluding one cluster, i11, with too few cells). All cross-validation designs were two-fold by evenly splitting data into training and testing in proportion to the original cluster sizes. All cross-validations were repeated 20 times each design.

Real data-guided simulations were used to mimic under-/over-partitioned scenarios (Figure 4). “Top nodes” under-partitions are cells merged into three broad classes: GABAergic inhibitory neurons, glutamatergic excitatory neurons, and neuroglial cells. “Mid nodes” under-partitions are cells merged into similar inhibitory neurons according to the constellation diagram of cluster network from the original study [28]; for the purpose of simulation, i1 and i5, i3 and i4, and i6, i8, and i9 were merged. For over-partitions, large cell clusters were split by running k-means clustering with k = 2 independently for each over-partitioned cluster. “Split e1” divided the excitatory cluster into two sub-clusters of sizes 180 and 119 cells, resulting in 16 (= 15 + 1) over-partitioned clusters. “Split i1, i2, i3” divided each of the inhibitory clusters into two sub-clusters of sizes 56 and 34, 39 and 38, 32 and 24 cells, respectively, resulting in 18 (= 15 + 3) over-partitioned clusters in total. NS-Forest marker genes were identified for each of the simulated datasets. Matching performances of the under-/over-partitioned datasets were evaluated through two-fold cross-validation repeated 20 times.

## Data availability

Two published single-nucleus RNA-seq datasets from the Allen Institute of Brain Science of human brain were used: i) cortical Layer 1 of middle temporal gyrus (MTG) [28] and ii) full thickness MTG [29] (https://portal.brain-map.org/atlases-and-data/rnaseq/human-mtg-smart-seq). The Layer 1 dataset contains expression data from 871 intact nuclei that form 16 cell type clusters, including four non-neuronal type clusters, one excitatory neuron type cluster, and 11 inhibitory neuron type clusters. The MTG dataset contains filtered expression data from 15,603 nuclei that form 75 cell type clusters, subdivided into six non-neuronal type clusters, 24 excitatory neuron type clusters, and 45 inhibitory neuron type clusters. These cell type clusters are regarded as transcriptionally distinct cell types with nomenclature asserted after iterative clustering analysis [13]. Gene-level read count values were preprocessed to log-CPM (counts per million) values for all nuclei.

The same high-level data processing steps were used for both datasets, although the details varied slightly:

1. Whole postmortem brain specimens or neurosurgical tissue samples were collected from adult male and female donors with ‘control’ condition (i.e. non-disease).
2. Nuclei were isolated from microdissected tissue pieces to avoid damage to neurons [44], and single nuclei were sorted using FACS instruments. The gating strategy included doublet detection gates and gates on neuronal marker NeuN signal.
3. RNA sequencing was performed using the SMART-Seq platform and multiplex library preparation.
4. STAR alignment of raw reads to human genome sequence, and sequence quantification using standard Bioconductor packages were performed. Gene expression levels were reported as counts per million (CPM) of exon and intron reads.
5. Nuclei passing quality control criteria were included for clustering analysis.
6. Iterative clustering procedure based on community detection were performed to group nuclei into transcriptomic cell types [13]. Dropouts were accounted for while selecting differentially expressed genes, and PCA was used for dimensionality reduction.
7. Clusters identified as donor-specific were flagged as outliers, and manually inspected for cluster-level QC before exclusion.

## Key Points

- Feature selection plays a key role in scRNAseq data integration of cell type clusters; using supervised feature selection instead of approaches based on dropout rates significantly improves the performance of existing cell type matching methods, e.g. ‘scmap’.
- The random forest-based ‘NS-Forest’ marker gene selection algorithm is an effective dimensionality reduction tool that produces an informative set of necessary and sufficient genes for characterizing reference cell types.
- The cluster-level cell type matching method ‘FR-Match’, which builds upon a non-parametric multivariate statistical test, shows robustness against missing reference cell types, i.e. novel query cell types.
- FR-Match precisely matched common cell types from two independent scRNAseq experiments that reflect the laminar characteristics of the two anatomically overlapping brain regions.
- FR-Match software provides barcode plots and minimum spanning tree graphs for the query and reference cell type clusters, which are user-friendly visualization tools for insightful data exploration of scRNAseq data clusters.

## Funding

The development and assessment of FR-Match was funded by the JCVI Innovation Fund, the Allen Institute for Brain Science, and the Chan–Zuckerberg Initiative DAF, an advised fund of the Silicon Valley Community Foundation (2018-182730). The funding bodies had no role in the design or conclusions of this study.

## Author contributions

Y.Z. and R.H.S. designed the study, conceived the statistical model, and wrote the manuscript. Y.Z. and B.D.A. developed the software suites. Y.Z. and B.D.A applied the software to real data analysis. Y.Z., B.D.A, T.E.B., J.A.M., and R.H.S. interpreted the real data analysis. R.D.H., T.E.B., J.A.M., R.H.S., and E.S.L. performed the single nucleus RNA sequencing experiments used. R.H.S. and E.S.L. supervised the work.

**Yun Zhang** is a Staff Scientist and Biostatistician in the Informatics Department at the J. Craig Venter Institute.

**Brian D. Aevermann** is Senior Bioinformatics Analyst in the Informatics Department at the J. Craig Venter Institute.

**Trygve E. Bakken** is an Assistant Investigator at the Allen Institute for Brain Science.

**Jeremy A. Miller** is a Senior Scientist at the Allen Institute for Brain Science. **Rebecca D. Hodge** is an Assistant Investigator at the Allen Institute for Brain Science. **Ed S. Lein** is a Senior Investigator at the Allen Institute for Brain Science.

**Richard H. Scheuermann** is a Professor and Director in the Informatics Department at the J. Craig Venter Institute.

## Notes

### Competing Interest Statement

The authors have declared no competing interest.

