## Supplementary Figure for "FR-Match: Robust matching of cell type clusters from single cell RNA sequencing data using the Friedman-Rafsky non-parametric test"

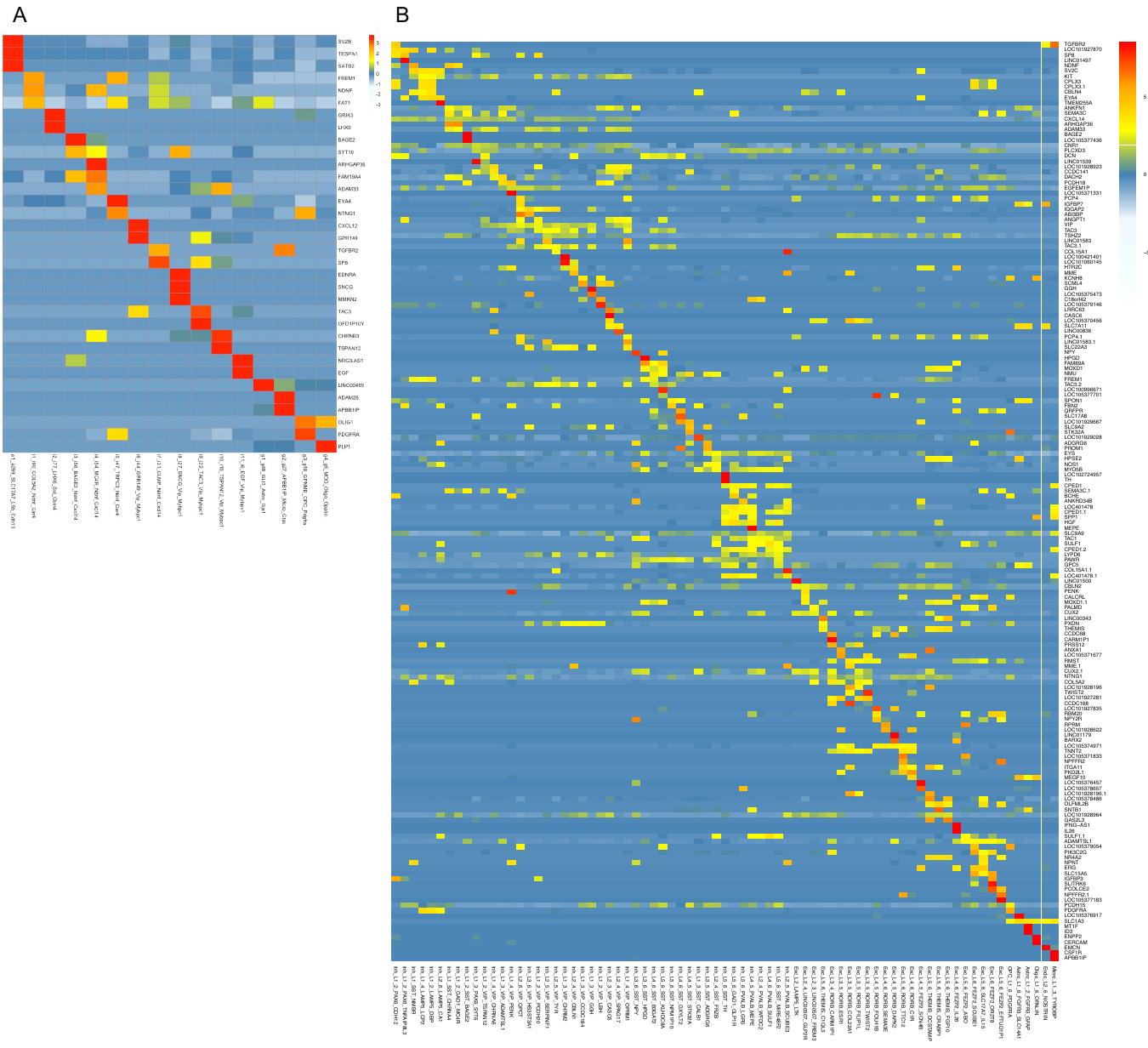

**Suppelementary Figure 1. Marker gene features selected by NS-Forest for cortical Layer 1 and full MTG datasets. (A)** Heatmap shows the median expression of NS-Forest marker genes for cell clusters in the Layer 1 dataset. There are 32 unique marker genes identified for the 16 Layer 1 cell type clusters providing a median F-beta classification score = 0.86. **(B)** Heatmap shows the median expression of NS-Forest marker genes for cell clusters in the full MTG dataset. There are 157 unique marker genes identified for the 75 full MTG cell type clusters providing a median F-beta classification score = 0.68.

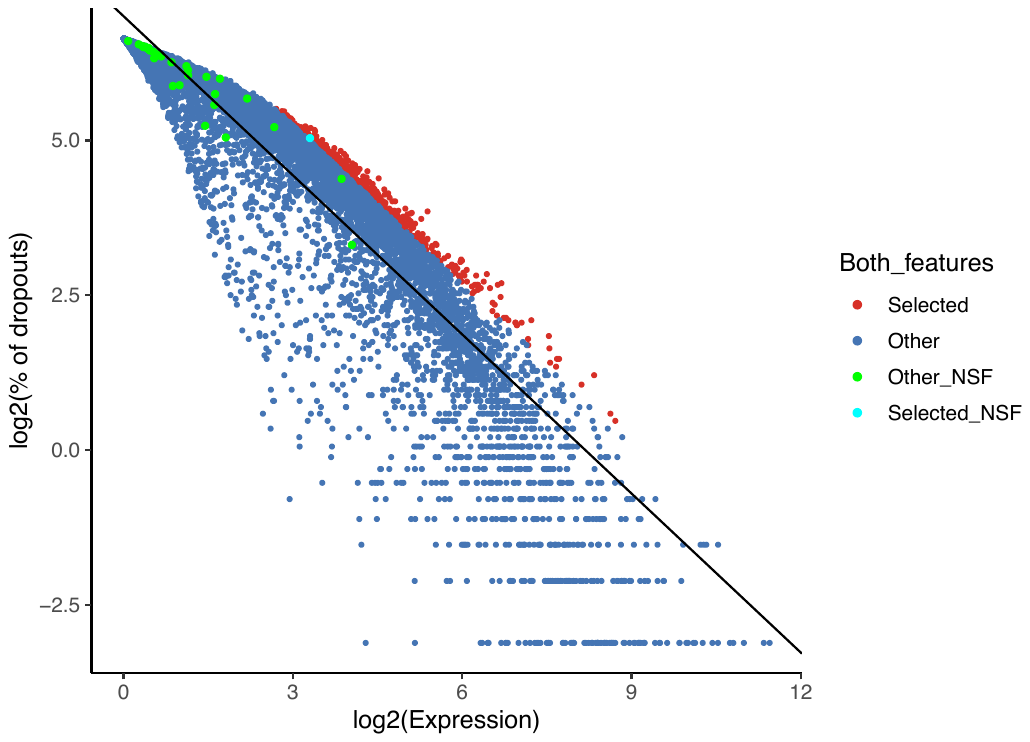

**Suppelementary Figure 2. scmap default selected features and NS-Forest marker genes in the expression-dropout-relationship plot used in scmap.** “Selected” points are 500 features selected by the default settings of scmap; “Other” are the genes not selected by scmap default; “Other_NSF” are the NS-Forest marker genes that overlap with the “Other” genes, “Selected_NSF” are NS-Forest marker genes that overlap with the “Selected”. There is only one gene that is selected by both scmap default and NS-Forest (“Selected_NSF”). Linear regression line was fitted for the gene expression and dropout rate relationship.

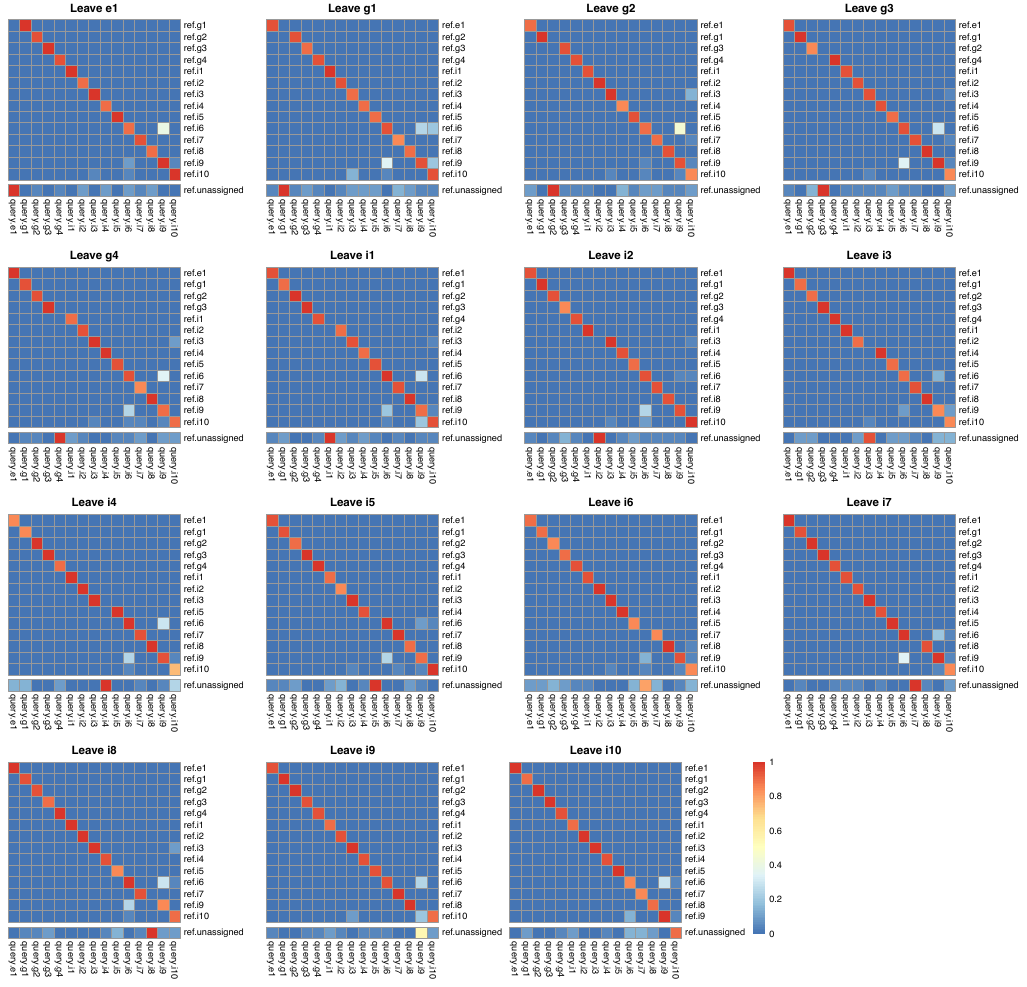

**Suppelementary Figure 3. Leave-one-cluster-out cross-validation results for FR-Match.** Heatmaps show the average matching results when each cell cluster was left out in turn using FR-Match.

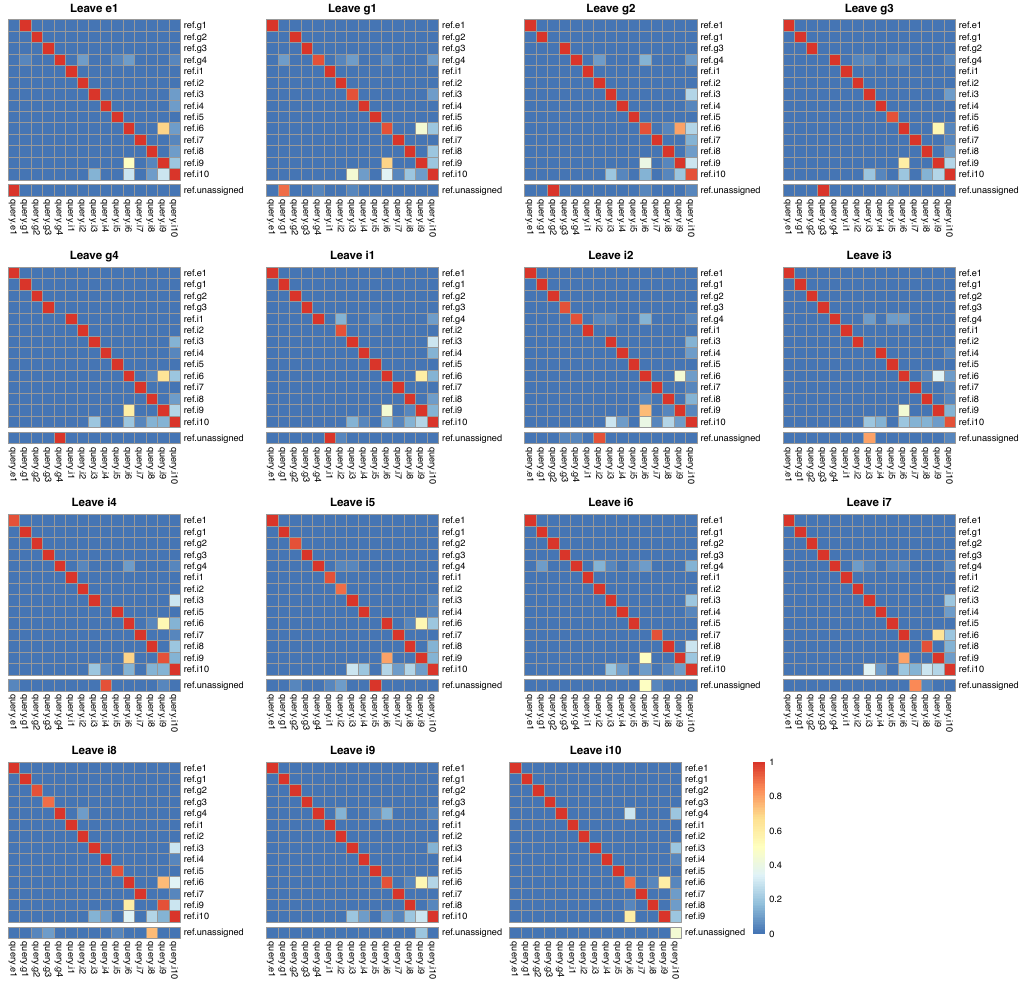

**Suppelementary Figure 4. Leave-one-cluster-out cross-validation results for FR-Match with p-value adjustment.** Heatmaps show the average matching results when each cell cluster was left out in turn using FR-Match with p-value adjustment.

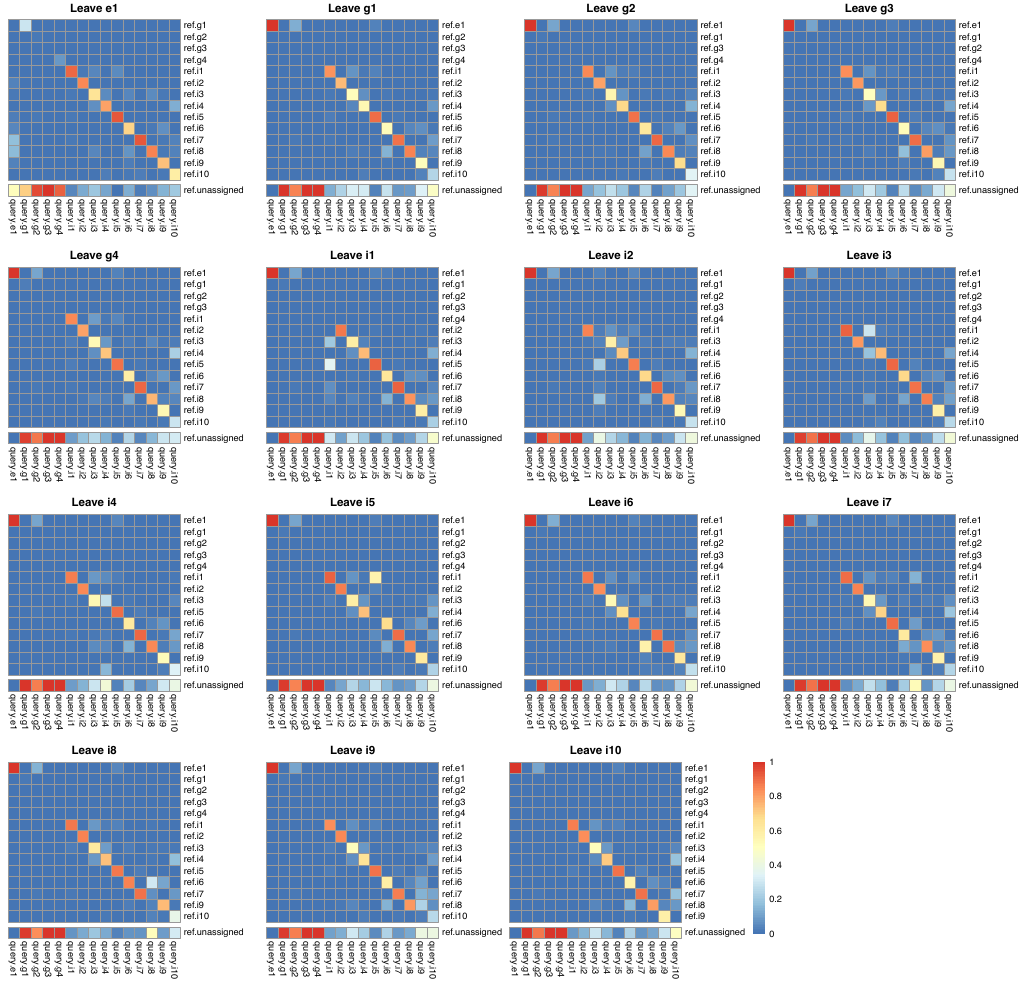

**Suppelementary Figure 5. Leave-one-cluster-out cross-validation results for scmap.** Heatmaps show the average matching results when each cell cluster was left out in turn using scmap.

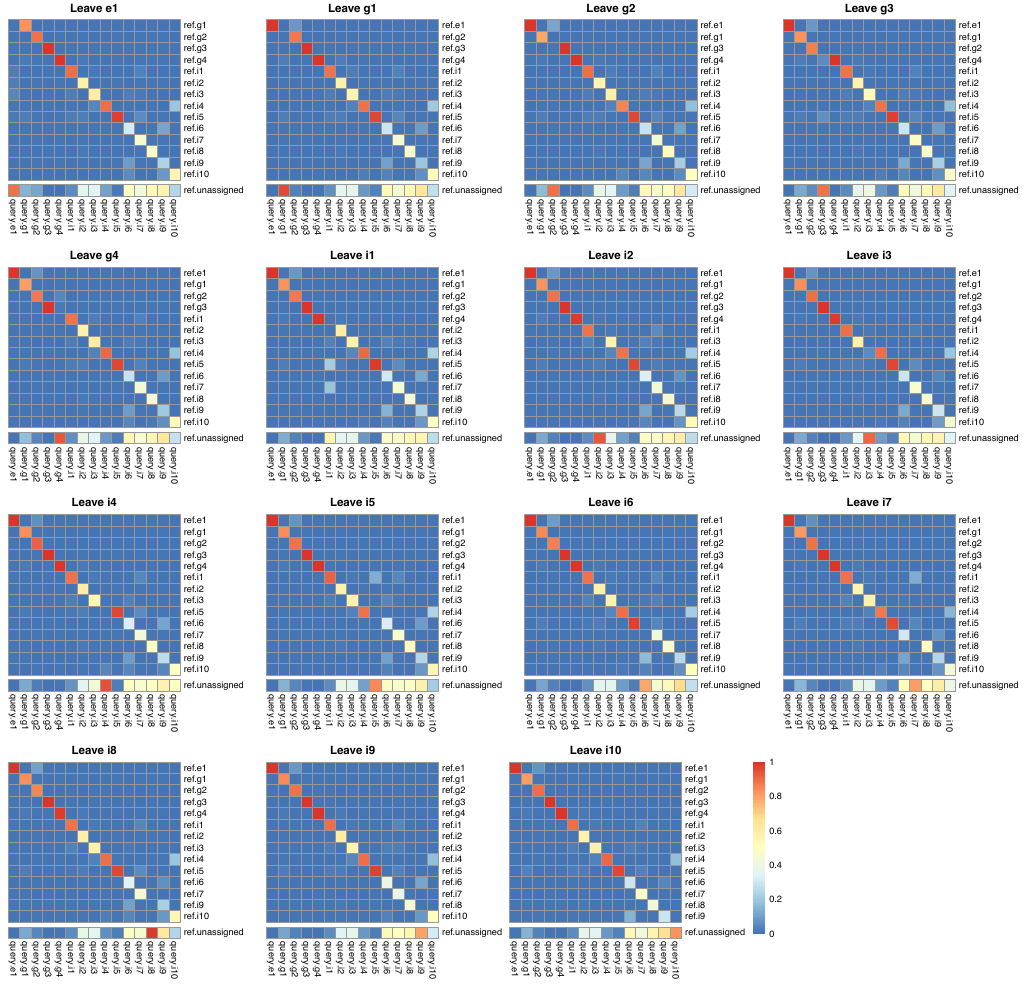

**Suppelementary Figure 6. Leave-one-cluster-out cross-validation results for scmap+NSF.** Heatmaps show the average matching results when each cell cluster was left out in turn using scmap with NSF markers.

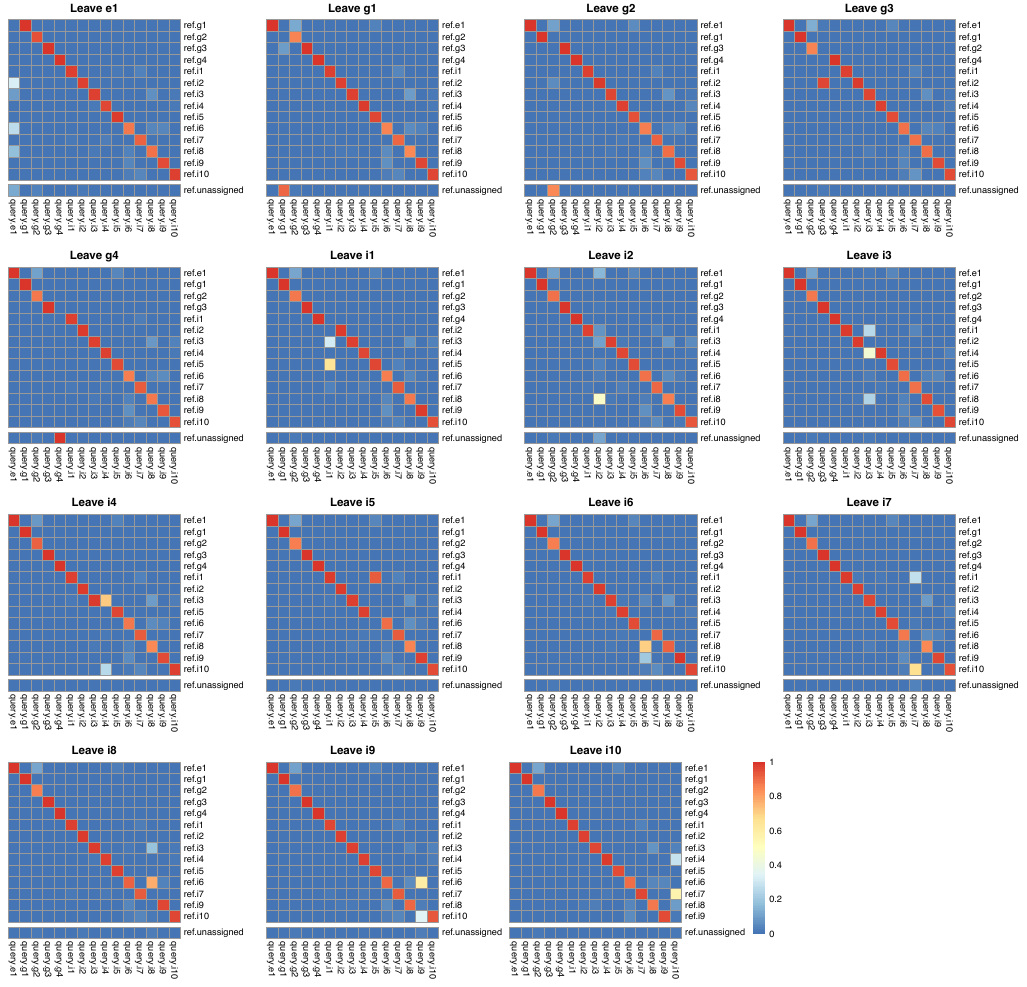

**Suppelementary Figure 7. Leave-one-cluster-out cross-validation results for scmap+NSF.ext.** Heatmaps show the average matching results when each cell cluster was left out in turn using scmap with the extended set of NSF markers.

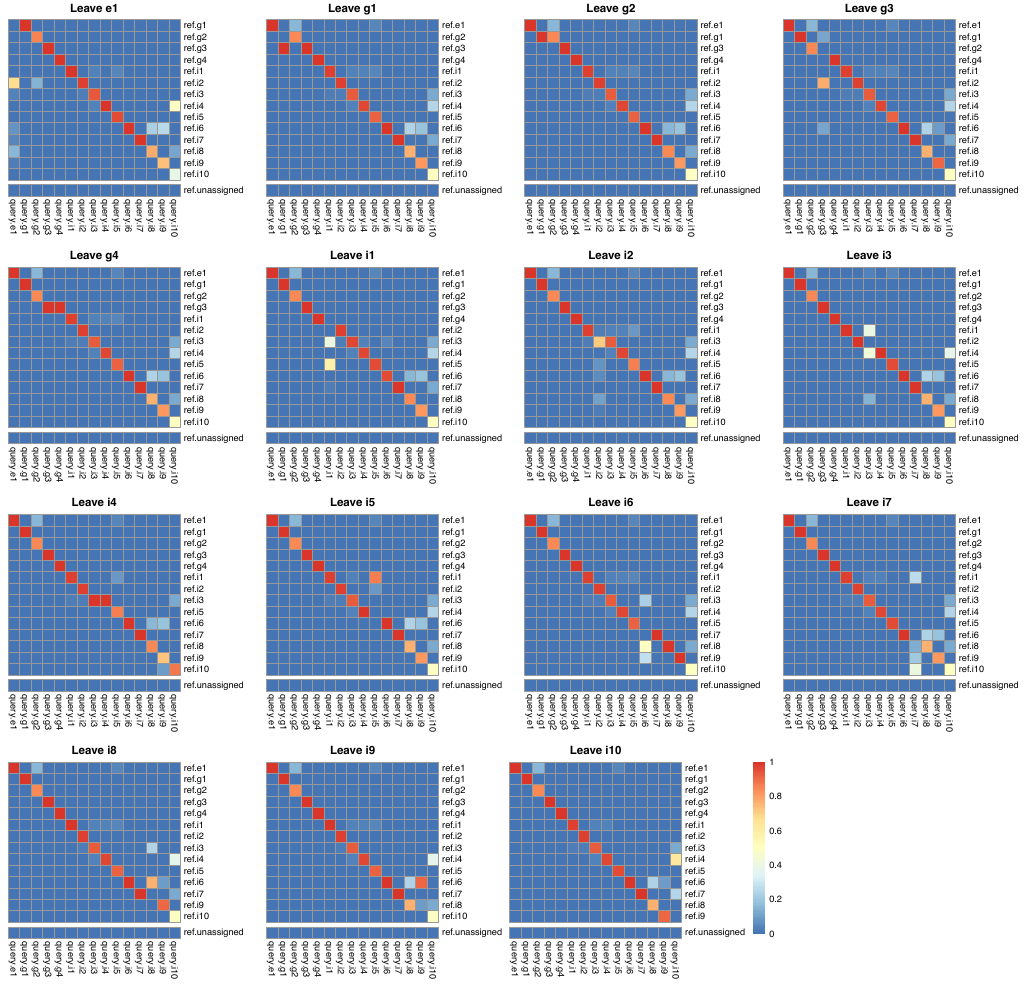

**Suppelementary Figure 8. Leave-one-cluster-out cross-validation results for Seurat.** Heatmaps show the average matching results when each cell cluster was left out in turn using Seurat.
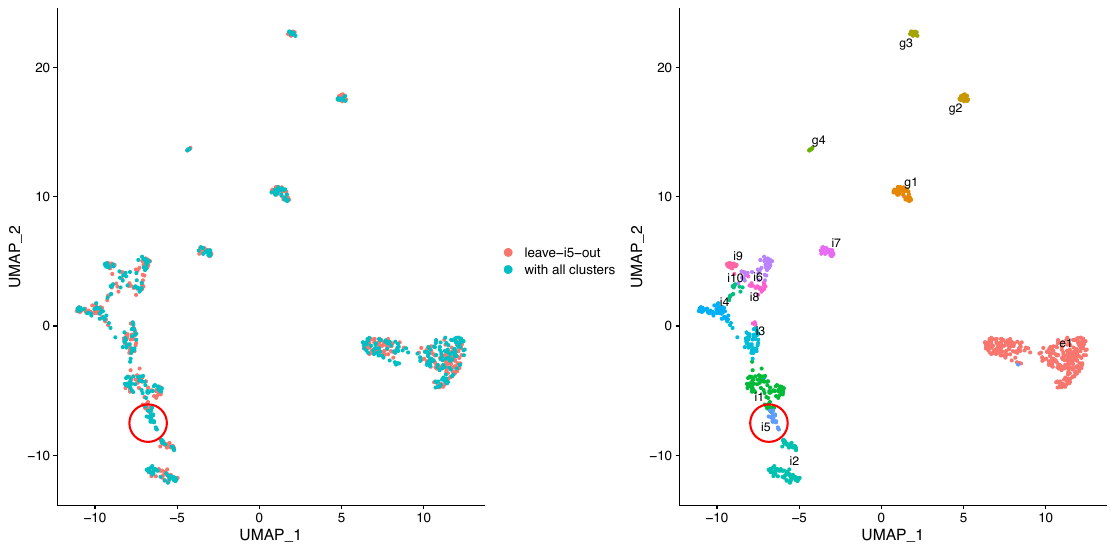

**Suppelementary Figure 9. UMAP of the leave-i5-out cross-validation datasets.** Left: Seurat integrated nuclei colored by reference (leave-i5-out) and query (with all clusters) datasets. Right: Seurat integrated nuclei colored by cell type labels. Red circle: query i5 nuclei.

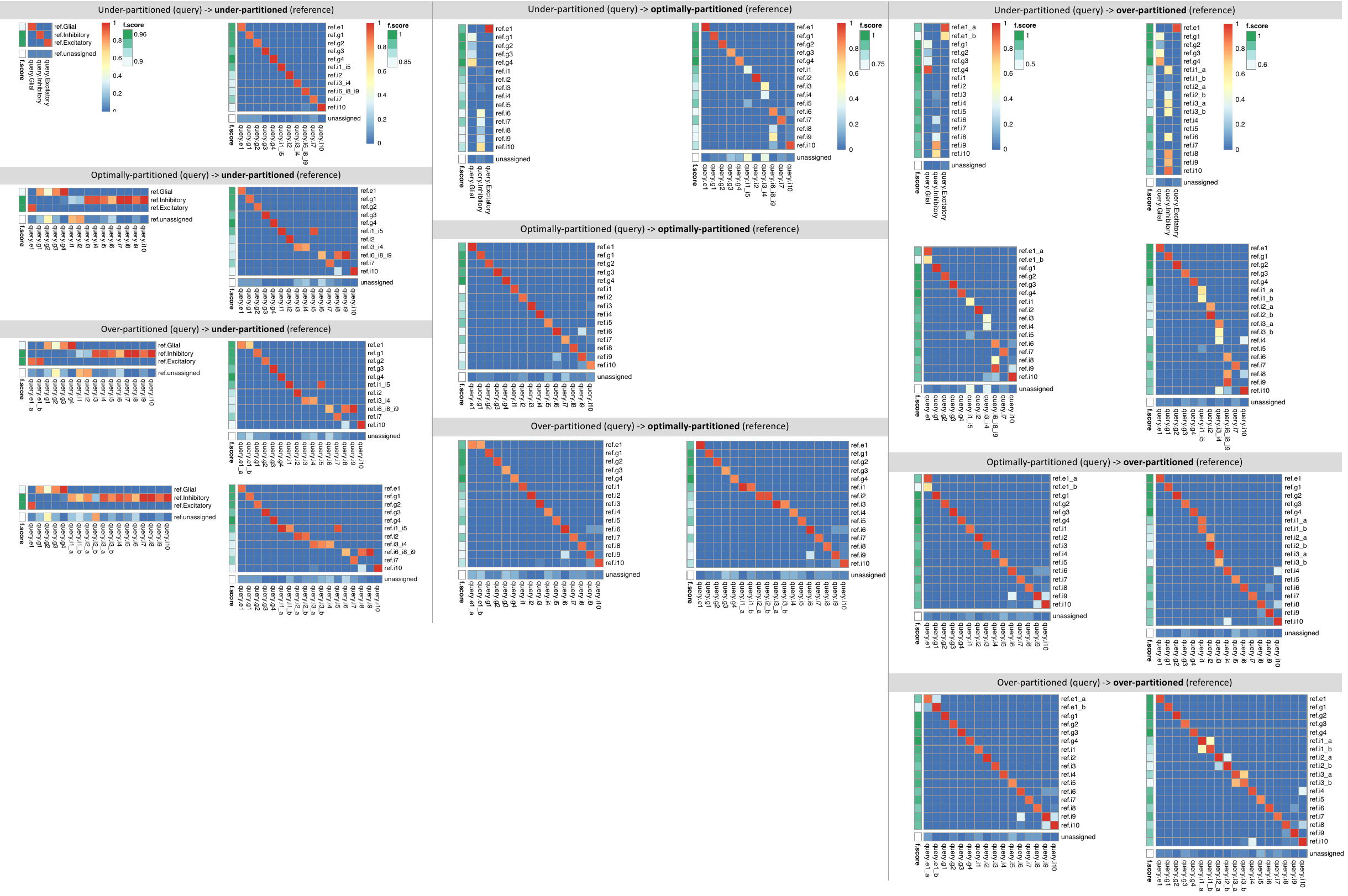

**Suppelementary Figure 10. FR-Match results for under-, optimally-, and over-partitioned clusters.** The optimally-partitioned clusters are the original clusters from the dataset. Base on the optimal partition, two under-partition scenarios were simulated by merging similar/hierarchically-connected clusters, and two over-partition scenarios were simulated by splitting the most abundant clusters (Figure 4a). Left: Matching results for under-partitioned reference clusters. Middle: Matching results for optimally-partitioned reference clusters. Right: Matching results for over-partitioned reference clusters.

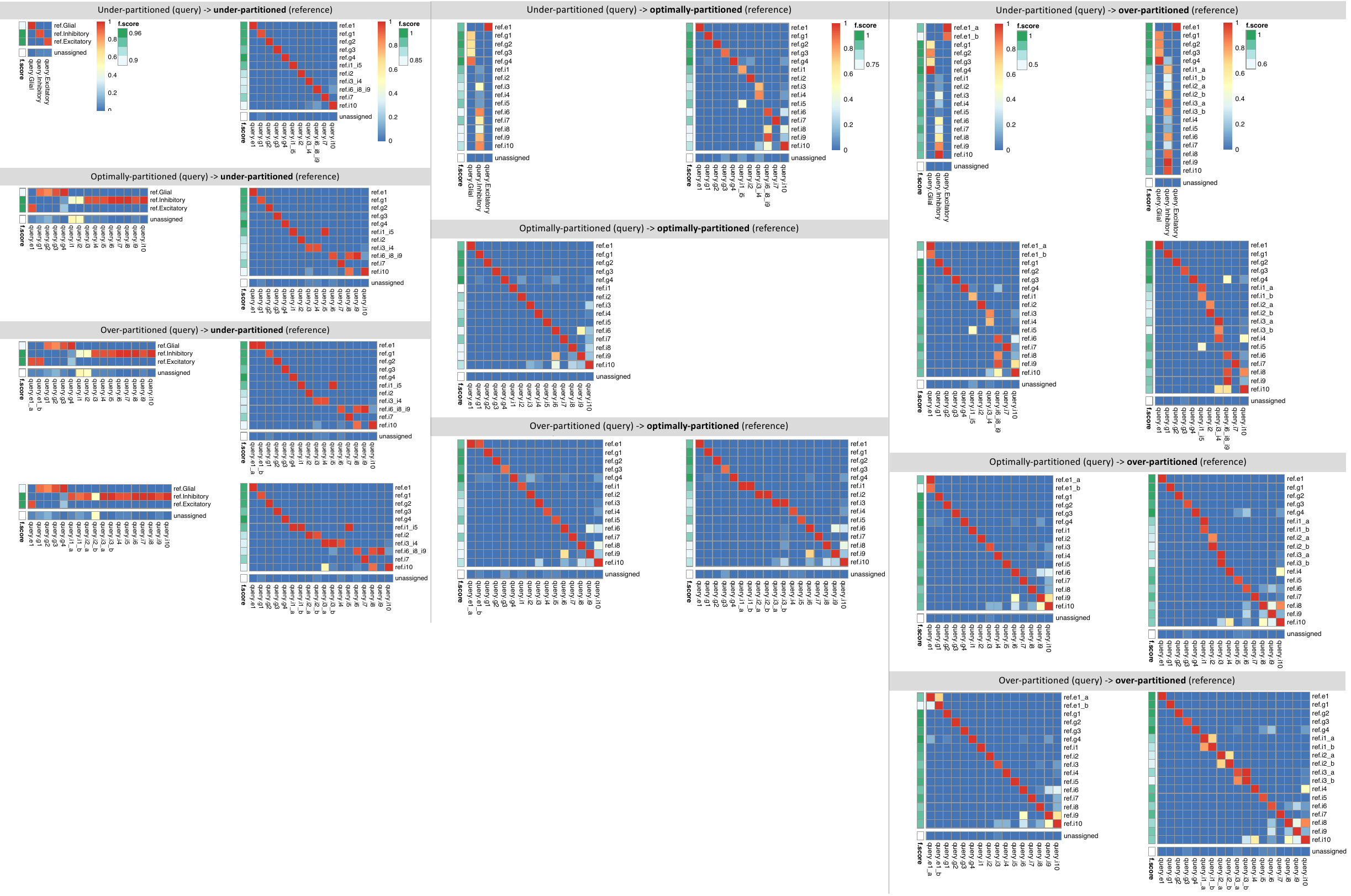

**Suppelementary Figure 11. FR-Match with p-value adjustment results for under-, optimally-, and over-partitioned clusters.** Left: Matching results for under-partitioned reference clusters. Middle: Matching results for optimally-partitioned reference clusters. Right: Matching results for over-partitioned reference clusters.

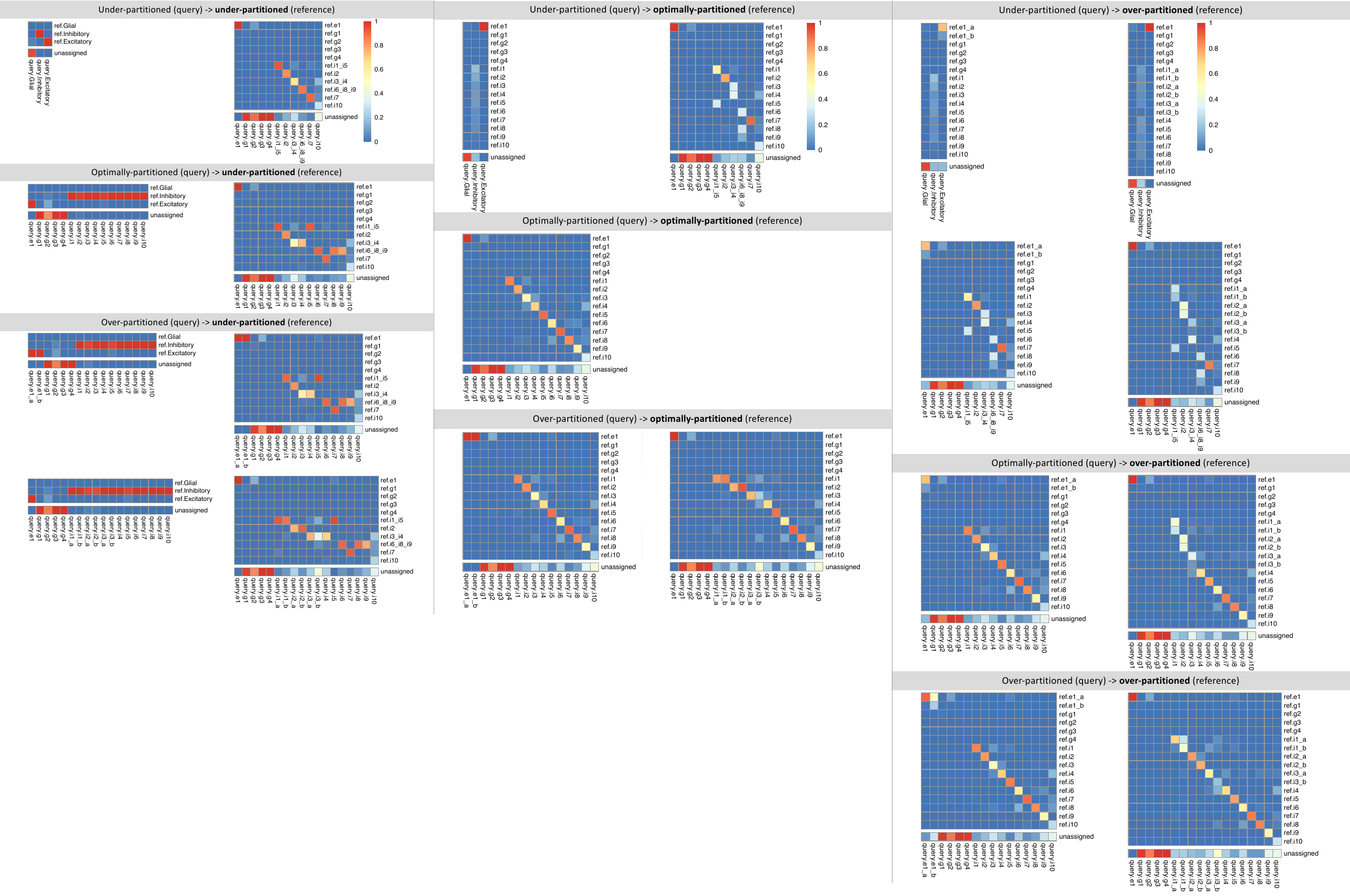

**Suppelementary Figure 12. Scmap results for under-, optimally-, and over-partitioned clusters.** Left: Matching results for under-partitioned reference clusters. Middle: Matching results for optimally-partitioned reference clusters. Right: Matching results for over-partitioned reference clusters.

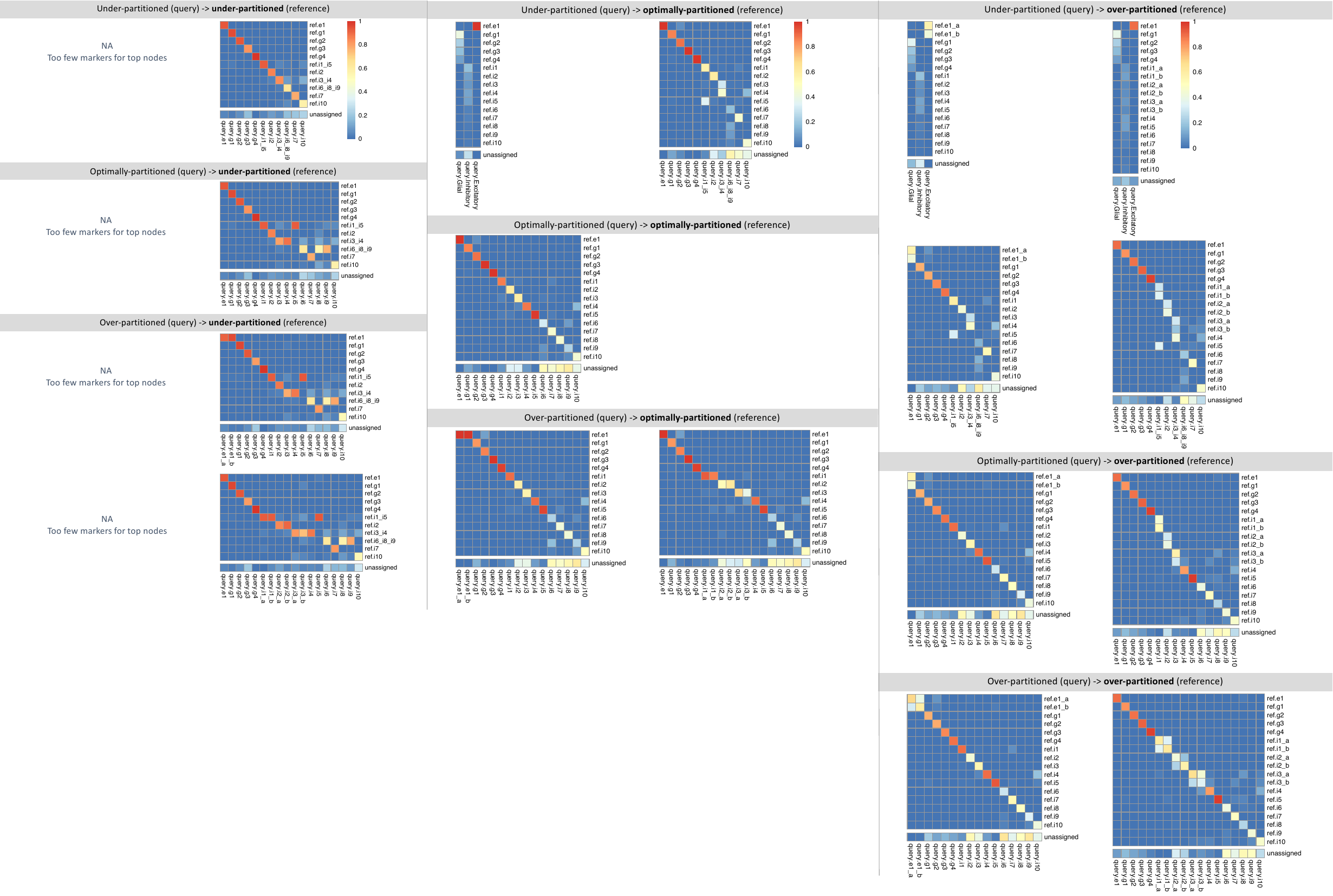

**Suppelementary Figure 13. Scmap+NSF results for under-, optimally-, and over-partitioned clusters.** Left: Matching results for under-partitioned reference clusters. Middle: Matching results for optimally-partitioned reference clusters. Right: Matching results for over-partitioned reference clusters.

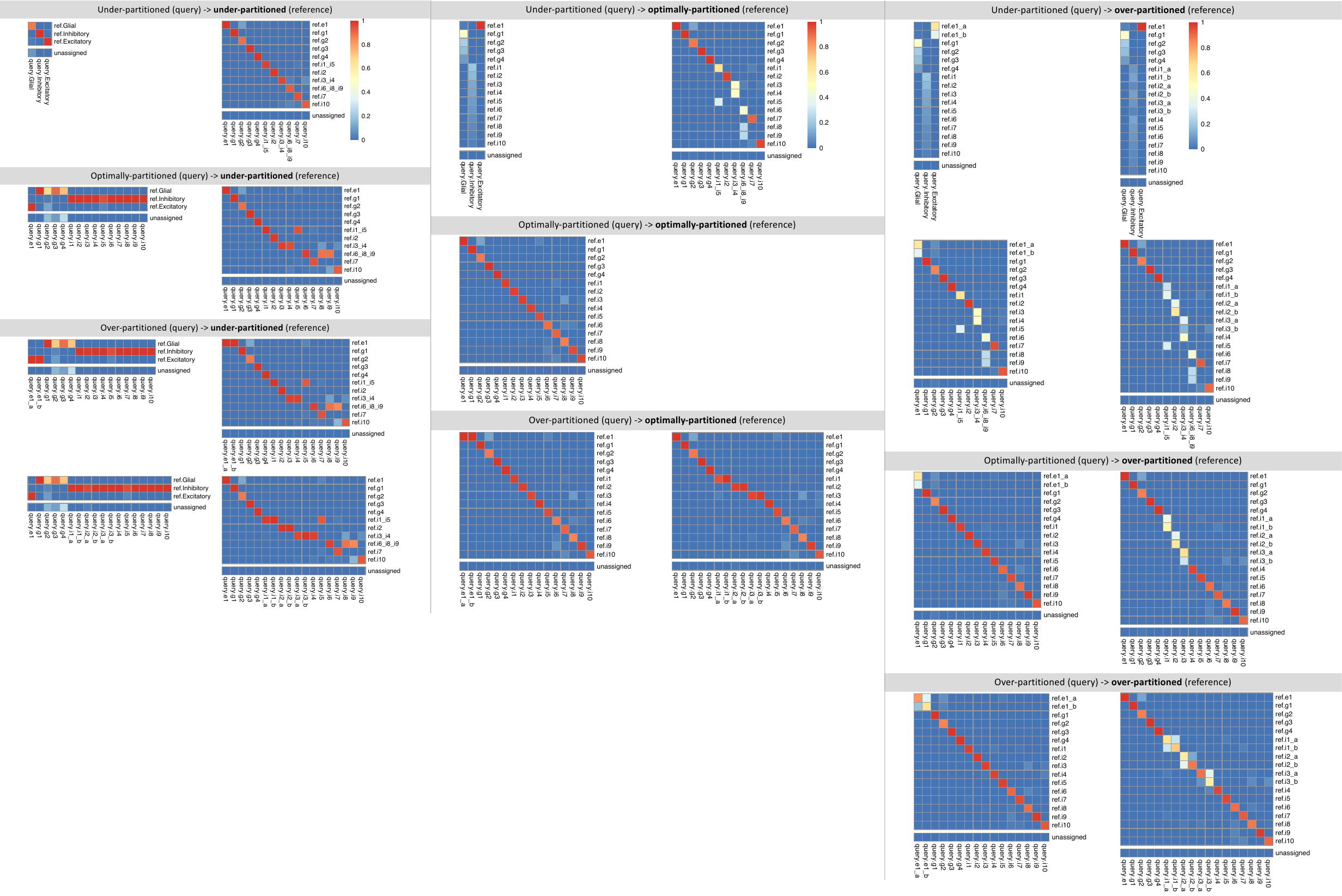

**Suppelementary Figure 14. Scmap+NSF.ext results for under-, optimally-, and over-partitioned clusters.** Left: Matching results for under-partitioned reference clusters. Middle: Matching results for optimally-partitioned reference clusters. Right: Matching results for over-partitioned reference clusters.

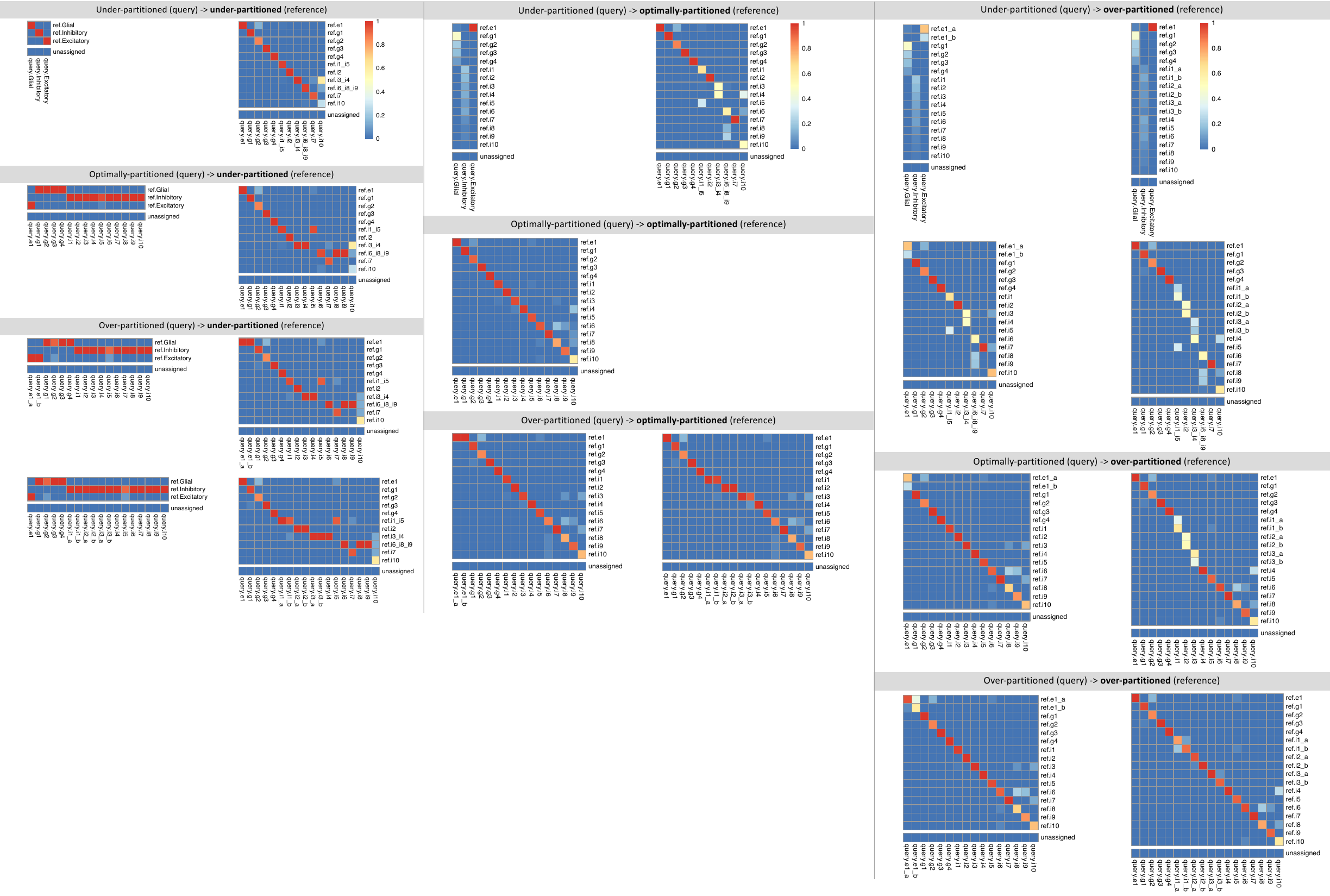

**Suppelementary Figure 15. Seurat results for under-, optimally-, and over-partitioned clusters.** Left: Matching results for under-partitioned reference clusters. Middle: Matching results for optimally-partitioned reference clusters. Right: Matching results for over-partitioned reference clusters.

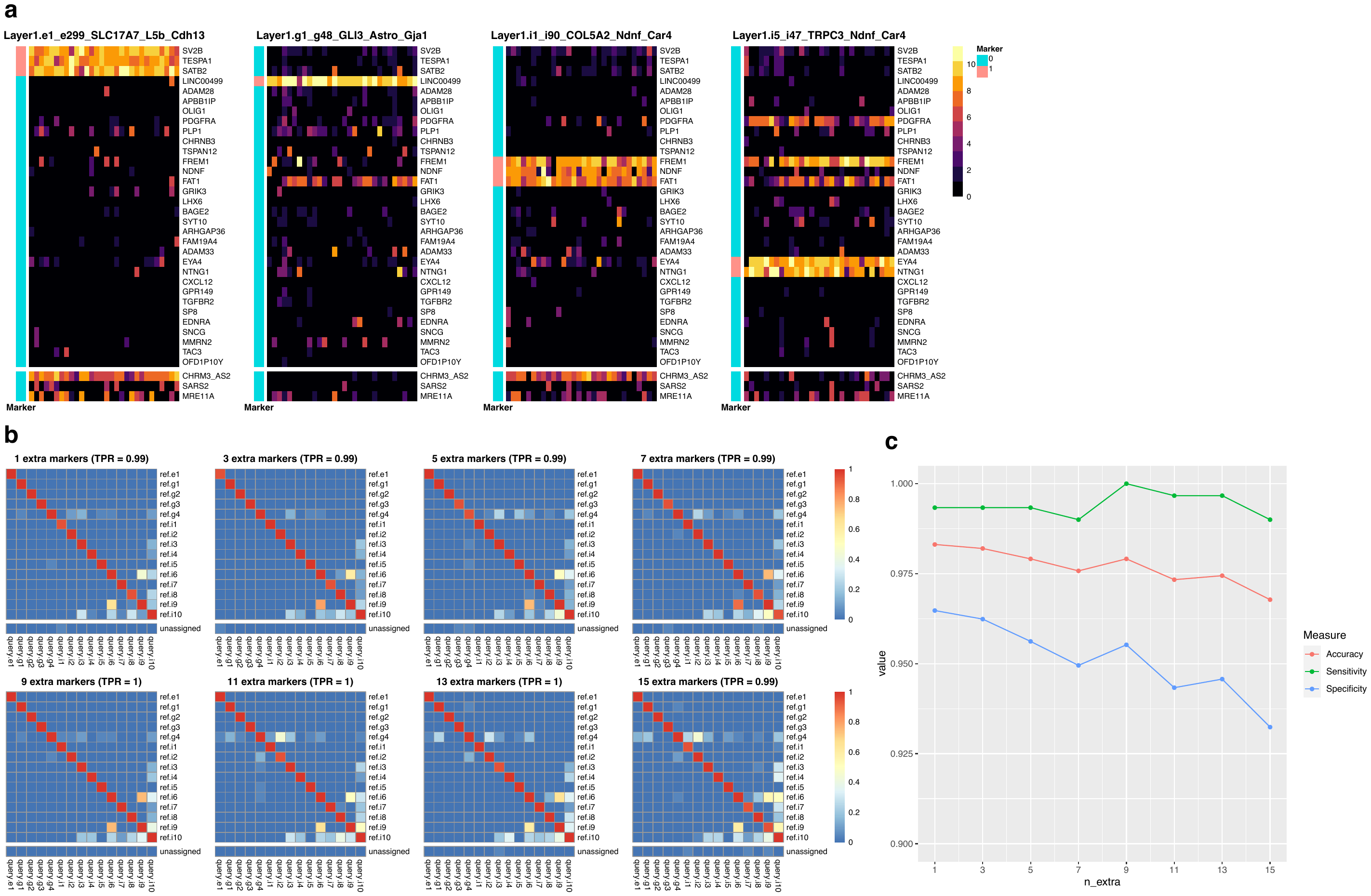

**Figure S16. Simulation for adding non-informative genes in the Layer 1 cell type matching analysis. (a)** Example “barcode” plots where 3 extra genes were randomly selected (bottom three rows). **(b)** Over 20 simulations, the average matching results for FR-Match adj. when adding 1-15 extra non-informative genes to the informative marker gene list. **(c)** Trend of accuracy, sensitivity, and specificity with respect to increasing number of extra non-informative genes.

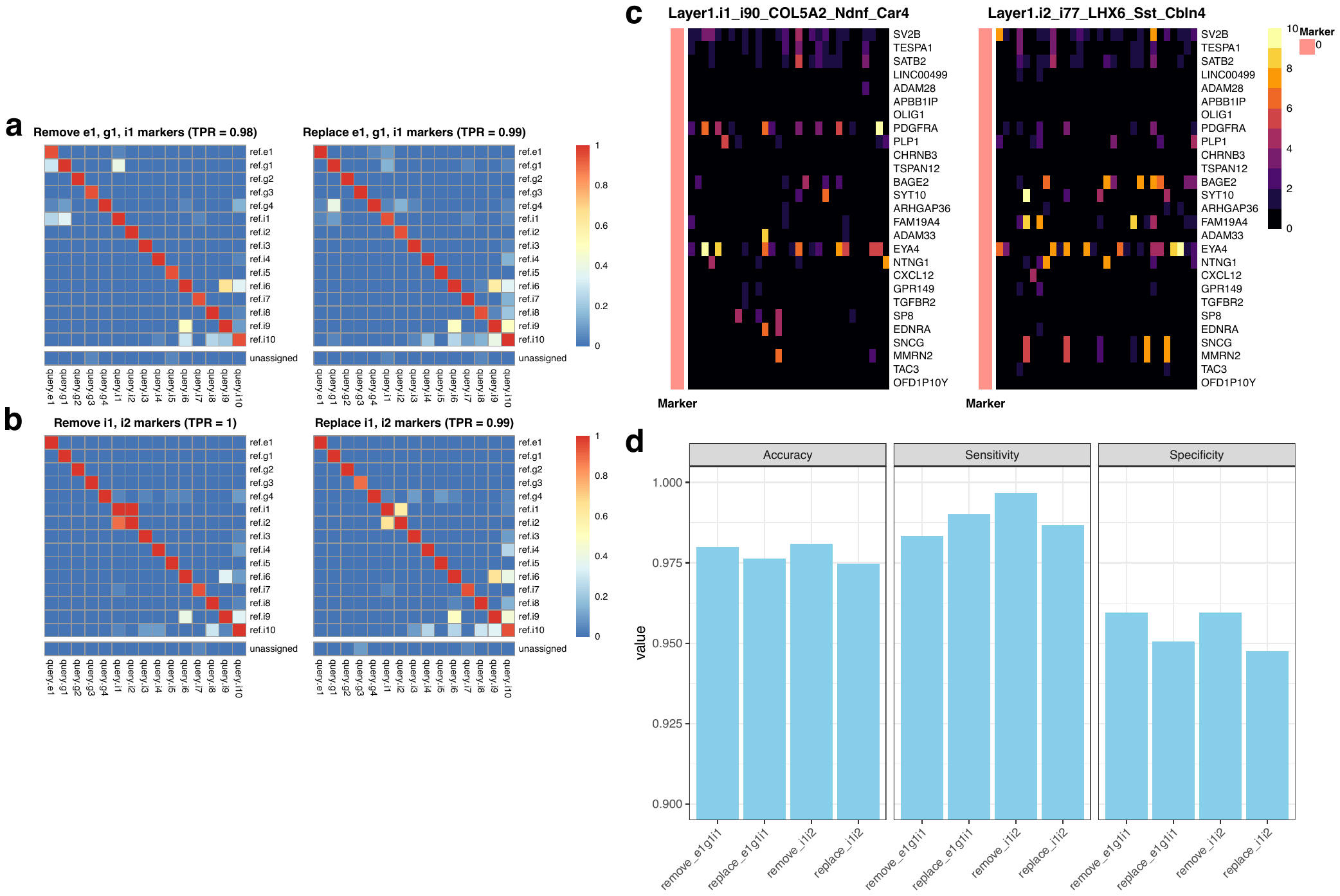

**Figure S17. Simulation for removing/replacing informative marker genes in the Layer 1 cell type matching analysis. (a)** Average matching results for FR-Match adj. when removing (left) or replacing (right) e1, g1, and i1 marker genes. **(b)** Average matching results for FR-Match adj. when removing (left) or replacing (right) i1 and i2 marker genes. **(c)** “Barcode” plots of cell types i1 and i2 where their marker genes are removed, leaving variable low signal. **(d)** Sensitivity, specificity, and accuracy for simulation cases in (a) and (b).

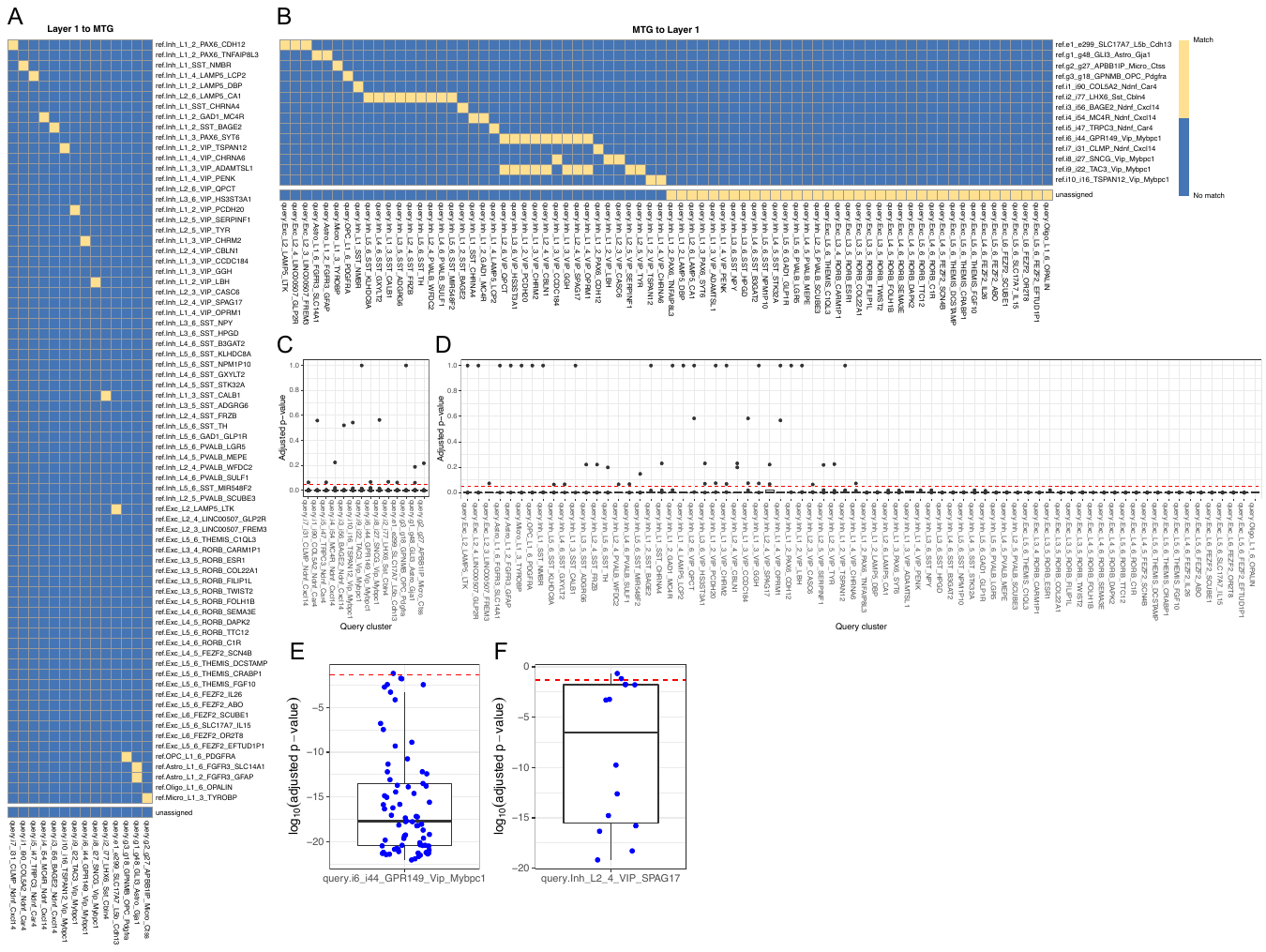

**Suppelementary Figure 18. FR-Match directional results and p-value distribution.** **(A)** One-way match results for directional matching from Layer 1 (query) to MTG (reference), which uses MTG marker genes. **(B)** One-way match results for directional matching from MTG (query) to Layer 1 (reference), which uses Layer 1 marker genes. **(C)** Box plots show the distribution of adjusted p-values for each Layer 1 query cluster. **(D)** Box plots show the distribution of adjusted p-values for each MTG query cluster. **(E)** A box plot from panel C in log10 scale. **(F)** A box plot from panel D in log10 scale. Red dash line in panels C-F shows the adjusted p-value cutoff = 0.05 for rejecting the null hypothesis that the clusters are matched.

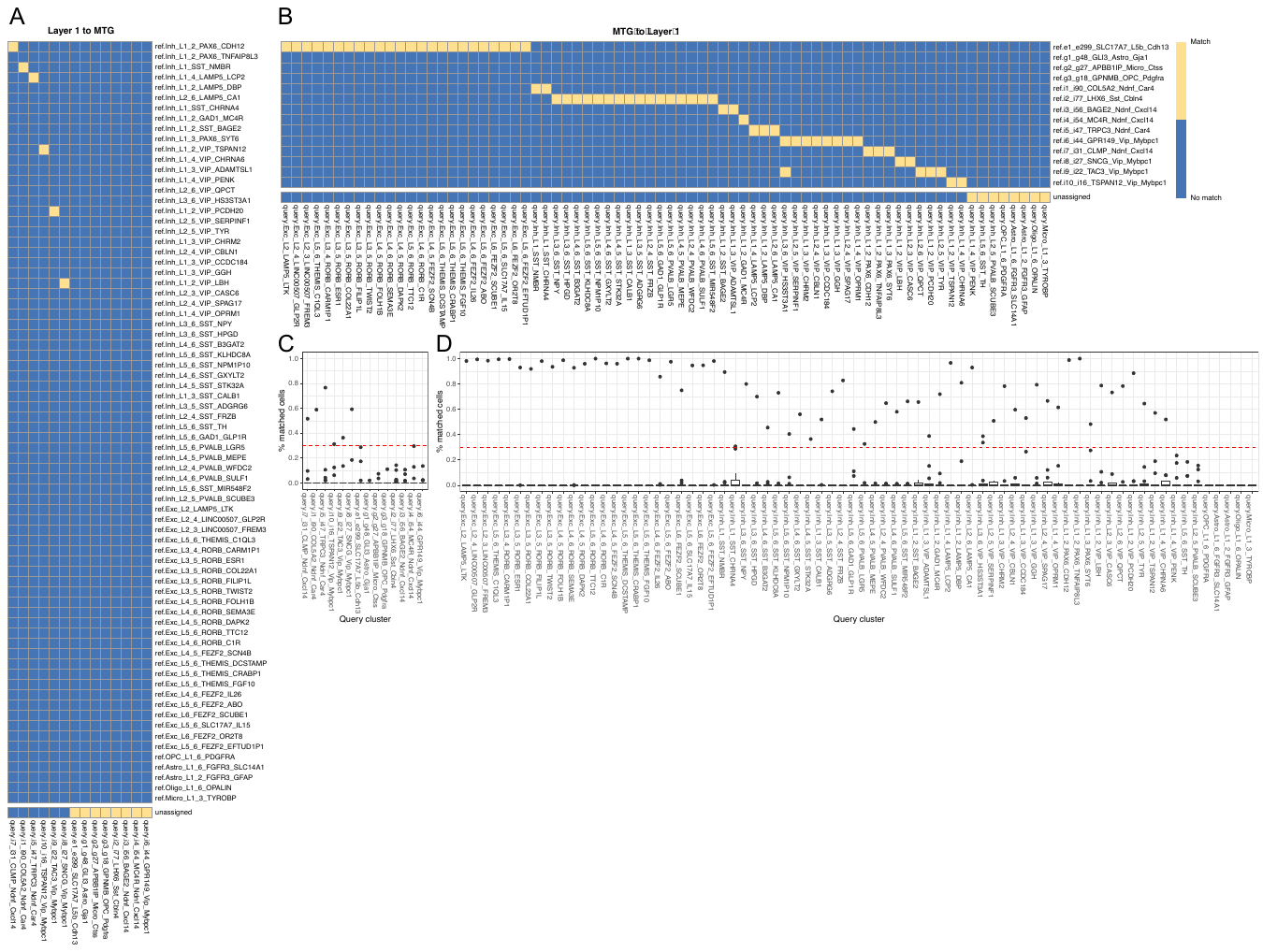

**Suppelementary Figure 19. Scmap directional matching results and p-value distributions.** **(A)** One-way matching results for directional matching from Layer 1 (query) to MTG (reference). **(B)** One-way matching results for directional matching from MTG (query) to Layer 1 (reference). **(C)** Box plots show the distribution of the cluster-level matching measure (% of matched cells) for each Layer 1 query cluster. **(D)** Box plots show the distribution of the cluster-level matching measure (% of matched cells) for each MTG query cluster. Red dash line in panels C-D shows the % of matched cells cutoff = 30%.

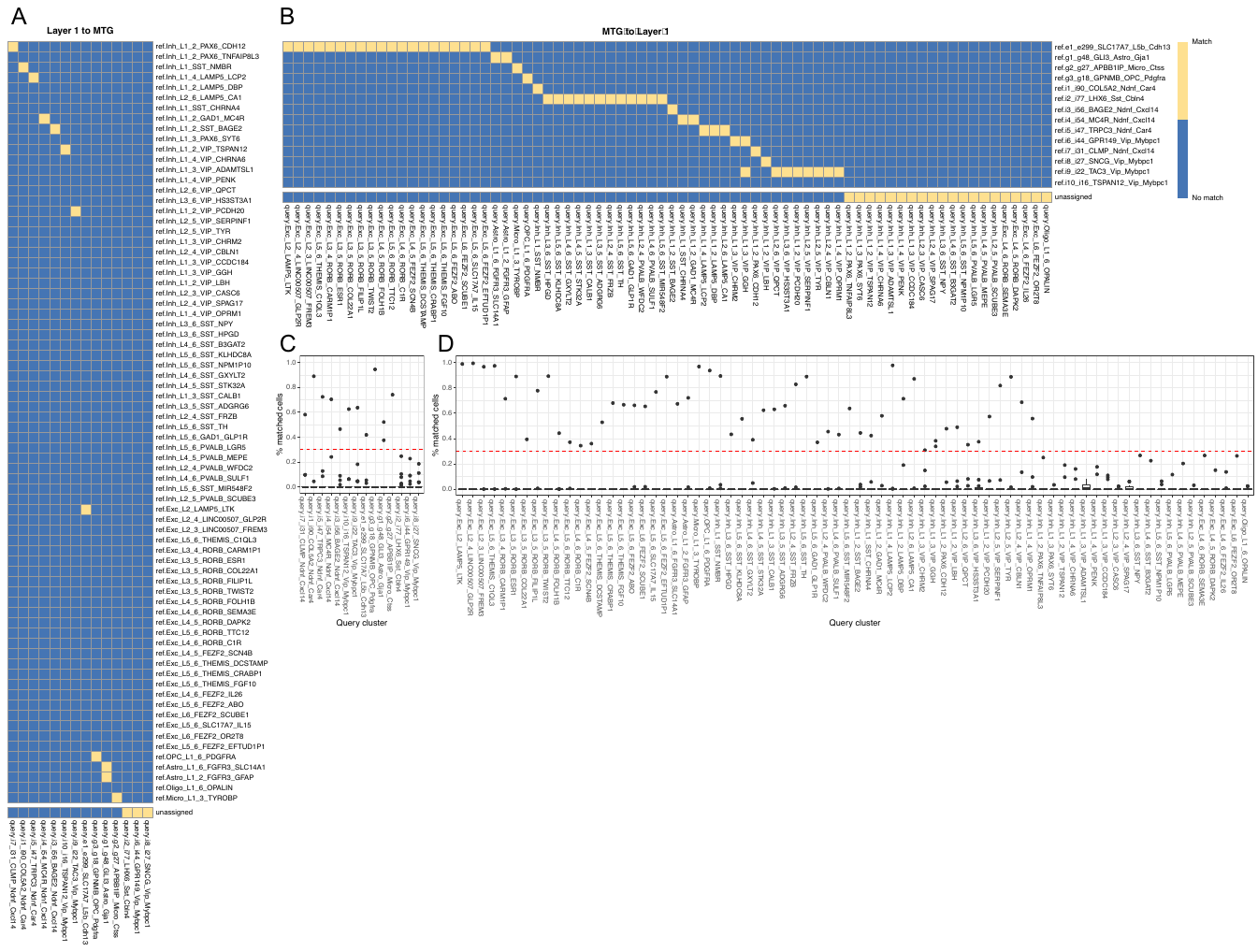

**Suppelementary Figure 20. Scmap+NSF directional matching results and p-value distributions.** **(A)** One-way matching results for directional matching from Layer 1 (query) to MTG (reference). **(B)** One-way matching results for directional matching from MTG (query) to Layer 1 (reference). **(C)** Box plots show the distribution of the cluster-level matching measure (% of matched cells) for each Layer 1 query cluster. **(D)** Box plots show the distribution of the cluster-level matching measure (% of matched cells) for each MTG query cluster. Red dash line in panels C-D shows the % of matched cells cutoff = 30%.

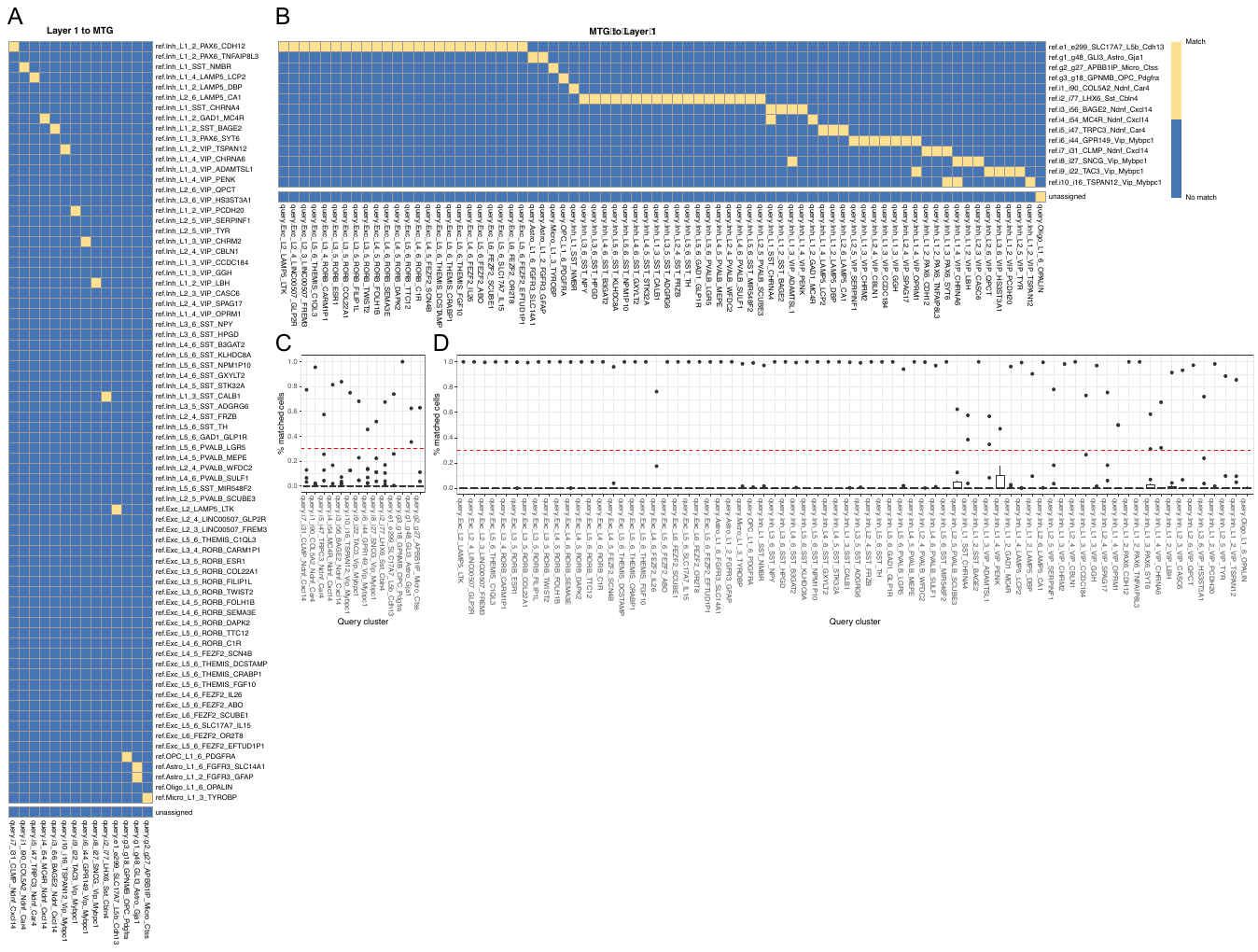

**Suppelementary Figure 21. Scmap+NSF.ext directional matching results and p-value distributions.** **(A)** One-way matching results for directional matching from Layer 1 (query) to MTG (reference). **(B)** One-way matching results for directional matching from MTG (query) to Layer 1 (reference). **(C)** Box plots show the distribution of the cluster-level matching measure (% of matched cells) for each Layer 1 query cluster. **(D)** Box plots show the distribution of the cluster-level matching measure (% of matched cells) for each MTG query cluster. Red dash line in panels C-D shows the % of matched cells cutoff = 30%.

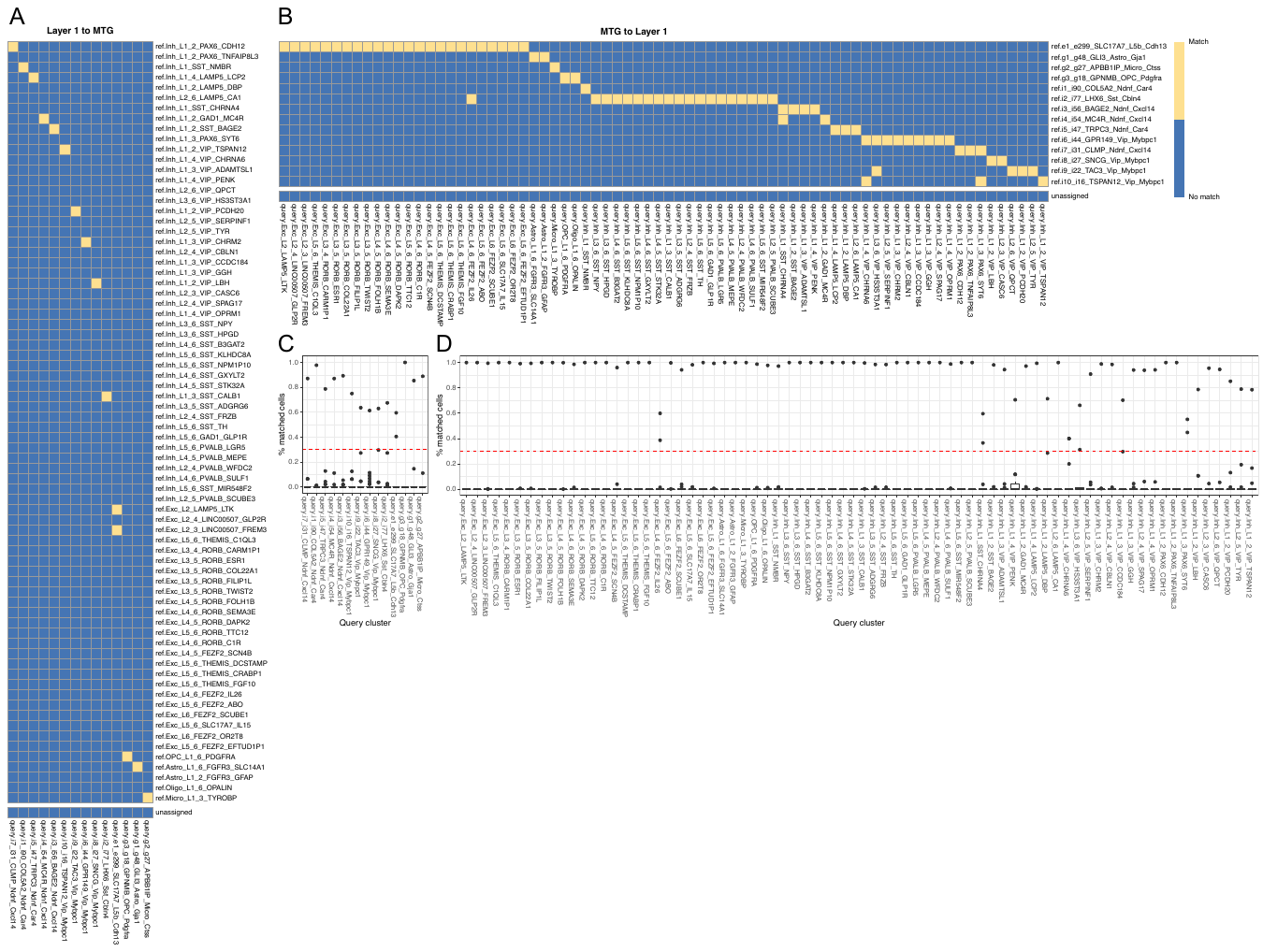

**Suppelementary Figure 22. Seurat directional matching results and p-value distributions.** **(A)** One-way matching results for directional matching from Layer 1 (query) to MTG (reference). **(B)** One-way matching results for directional matching from MTG (query) to Layer 1 (reference). **(C)** Box plots show the distribution of the cluster-level matching measure (% of matched cells) for each Layer 1 query cluster. **(D)** Box plots show the distribution of the cluster-level matching measure (% of matched cells) for each MTG query cluster. Red dash line in panels C-D shows the % of matched cells cutoff = 30%.

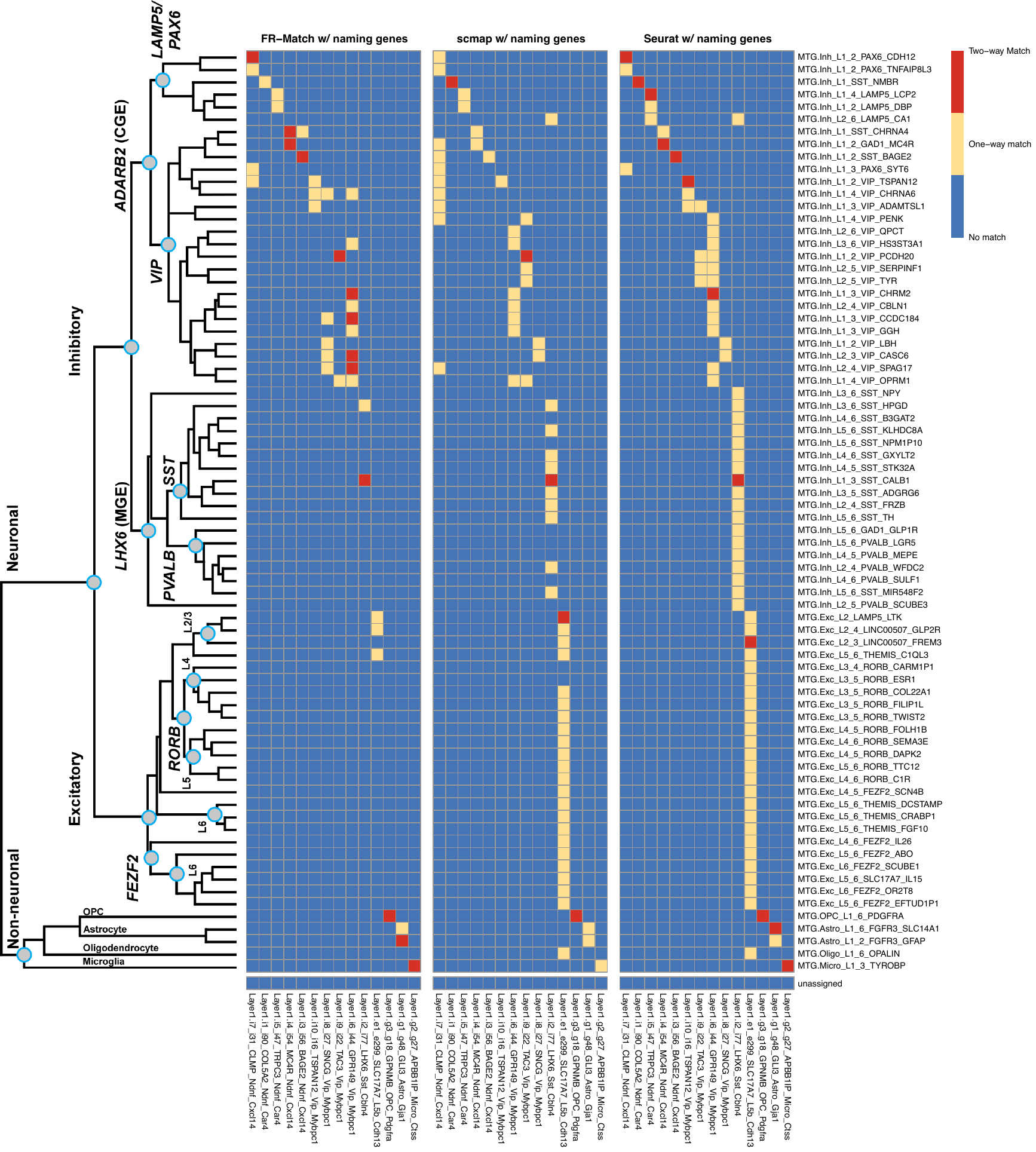

**Figure S23. Matching performance of FR-Match, scmap, and Seurat using the cell type naming genes of the Layer 1 and MTG cell types as an alternative feature selection approach.**

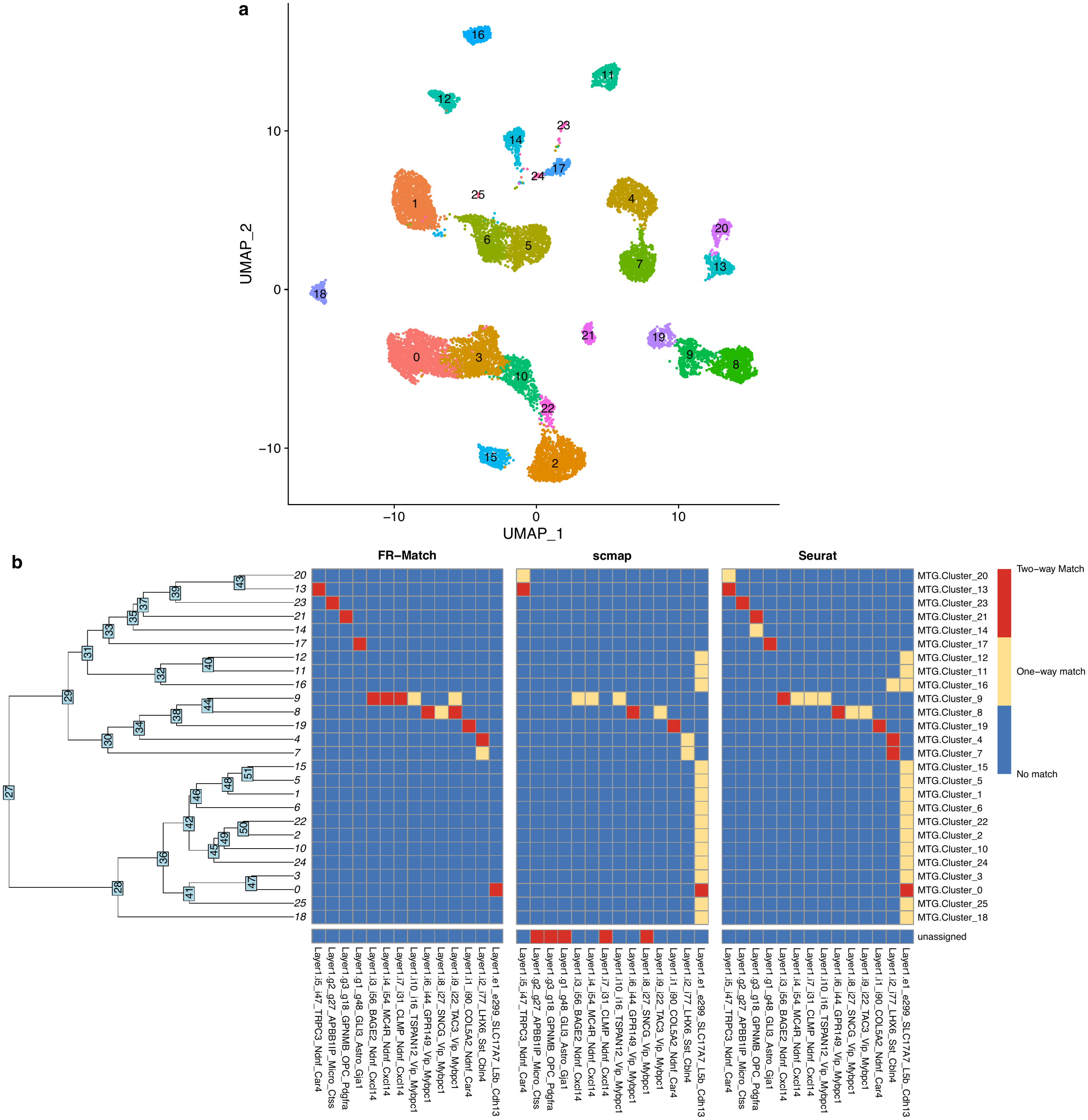

**Figure S24. Matching performance with Louvain clusters (implemented by the Seurat R package) of the MTG dataset. (a)** UMAP of the 26 Louvain clusters from the full MTG dataset. **(b)** Matching results of FR-Match, scmap, and Seurat with the Louvain clusters.

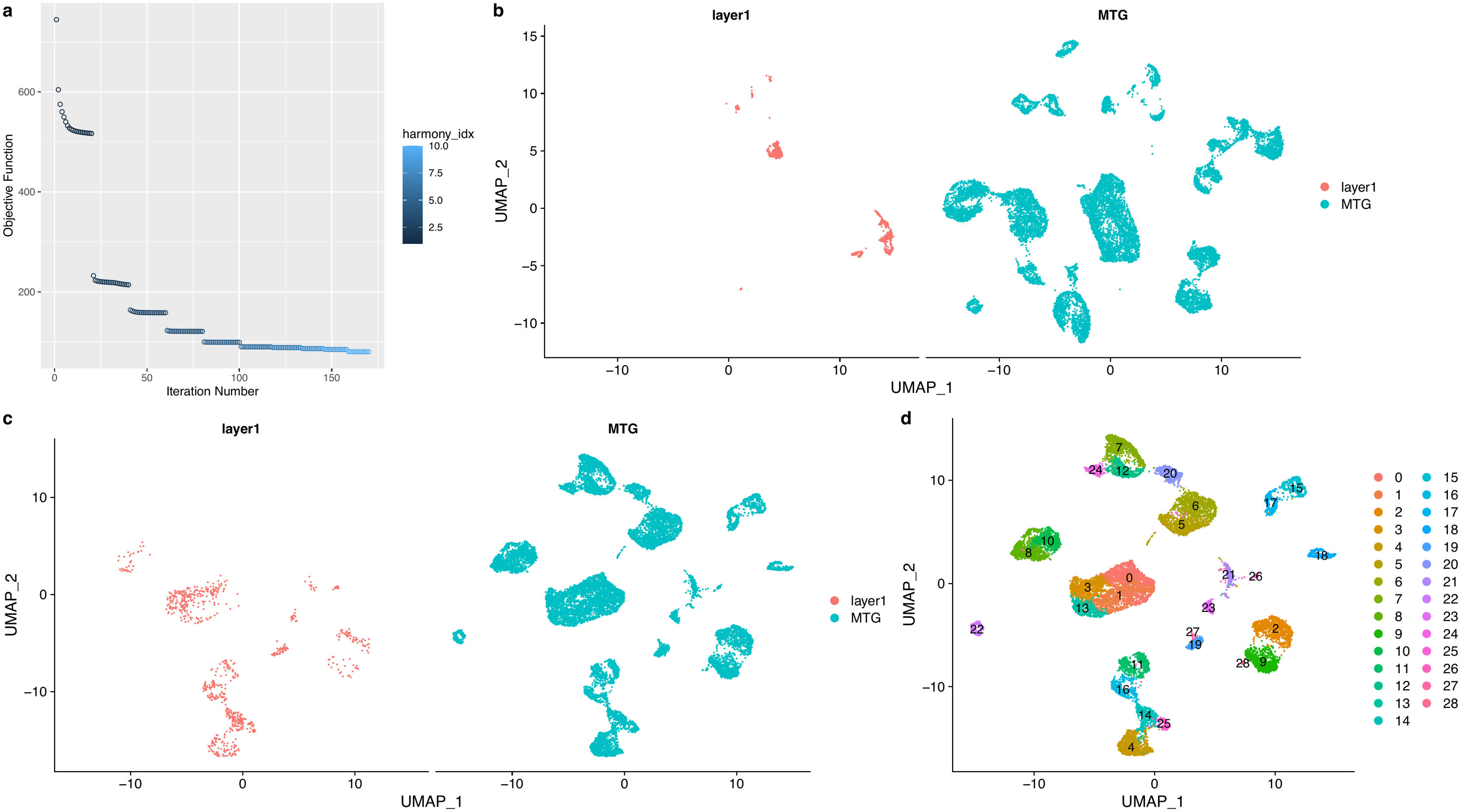

**Figure S25. Harmony batch integration. (a)** Convergence plot of the Harmony objective function. **(b)** UMAP embedding plot using the raw data. **(c)**  UMAP embedding plot of the Harmony integrated data. **(d)** Joint Louvain clustering (resolution = 1) on the Harmony integrated data.

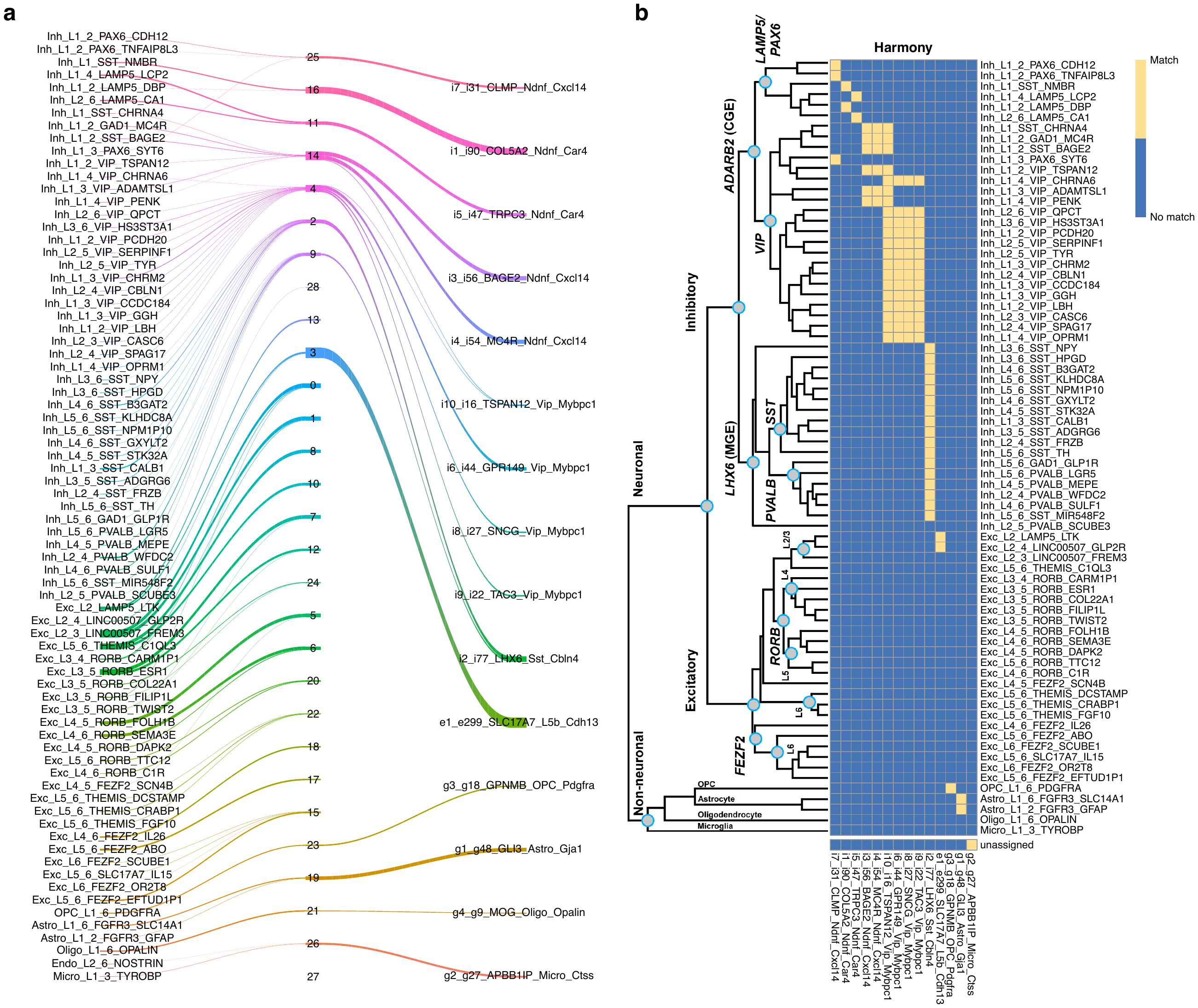

**Figure S26. Harmony river plot and inferred cell type matches. (a)** River plot visualization of the known cell types from individual datasets (left-full MTG and right-Layer 1 columns) through the joint cluster membership (middle column). **(b)** The cell type matching heatmap inferred from the edges of the river plot, with a match indicating that there exists a path between the two cell types on the river plot.

**Figure S27. LIGER batch integration. (a)** LIGER selects highly variable genes from both datasets in the data preprocessing step. Non-negative matrix factorization is performed on the subspace of the pooled variable genes from both datasets. **(b)** TSNE embedding plot using the raw data. **(c)**  TSNE embedding plot of the LIGER integrated data. **(d)** Joint Louvain clustering (resolution = 1) on the LIGER integrated data.

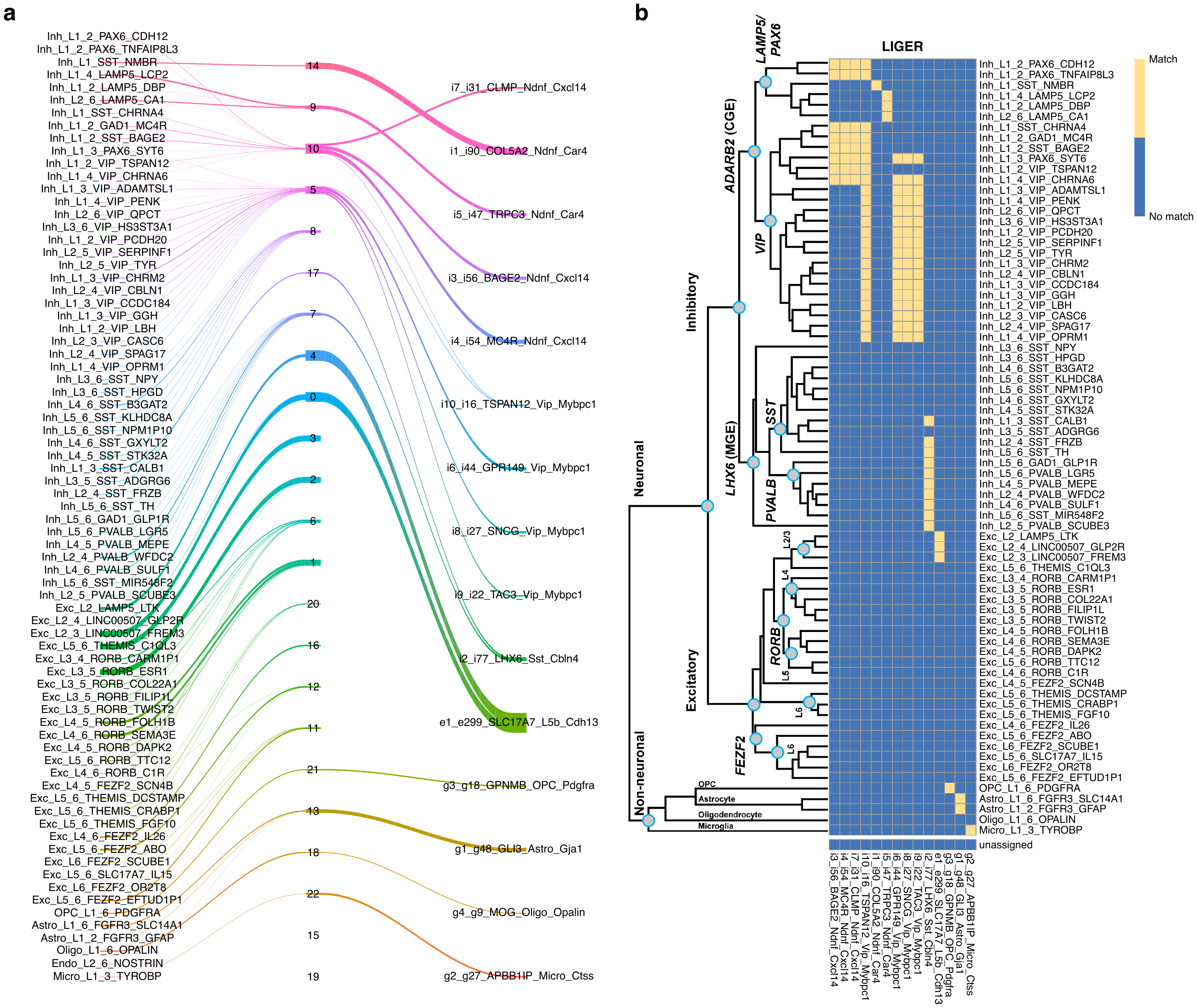

**Figure S28. LIGER river plot and inferred cell type matches. (a)** River plot visualization of the known cell types from individual datasets (left-full MTG and right-Layer 1 columns) through the joint cluster membership (middle column). **(b)** The cell type matching heatmap inferred from the edges of the river plot, with a match indicating that there exists a path between the two cell types on the river plot.

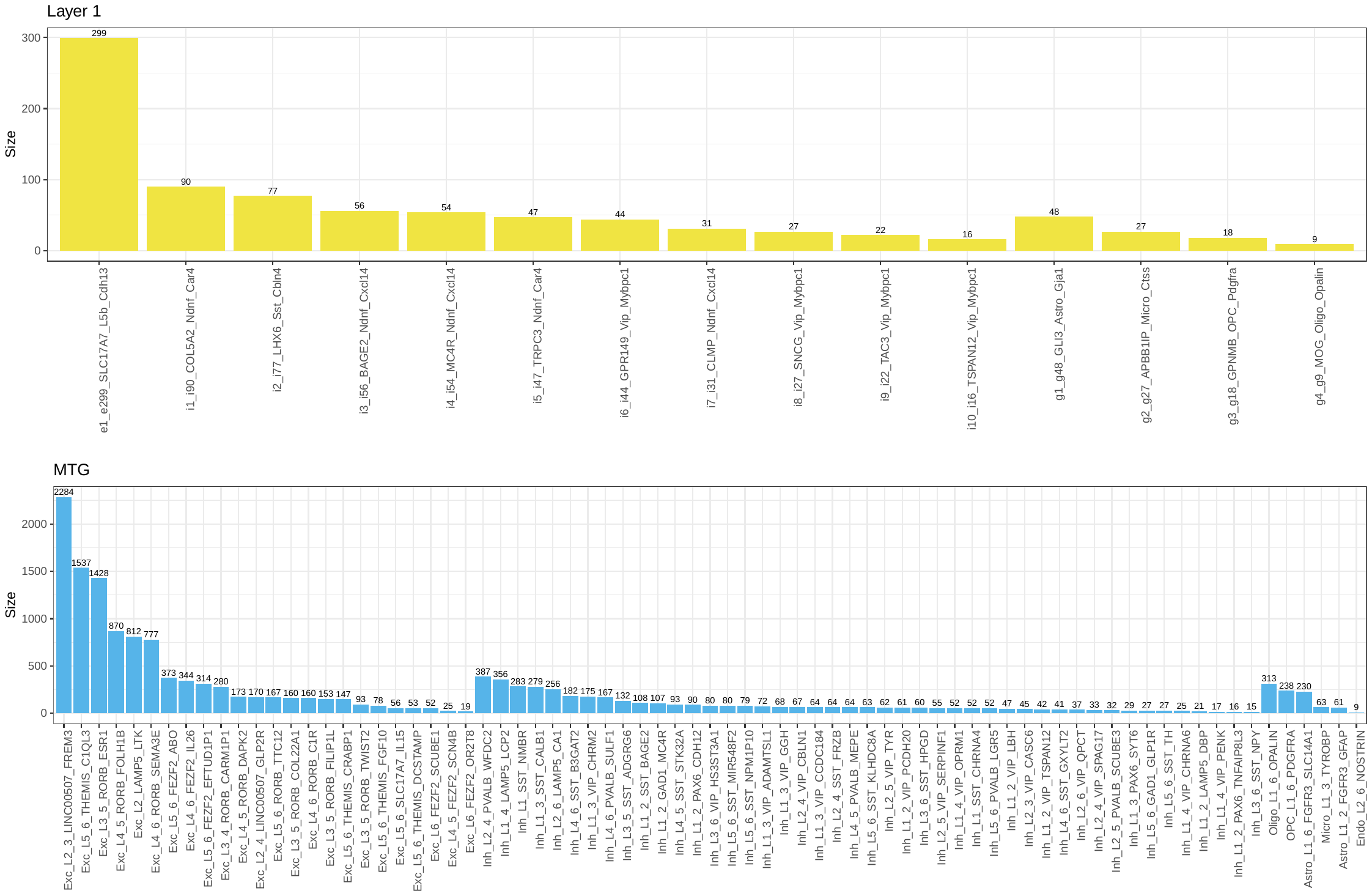

**Suppelementary Figure 29. Cluster sizes in Layer 1 and full MTG datasets.** Clusters are ordered by excitatory clusters ranked by size (number of nuclei per cluster), inhibitory clusters ranked by size, and non-neuronal clusters ranked by size.

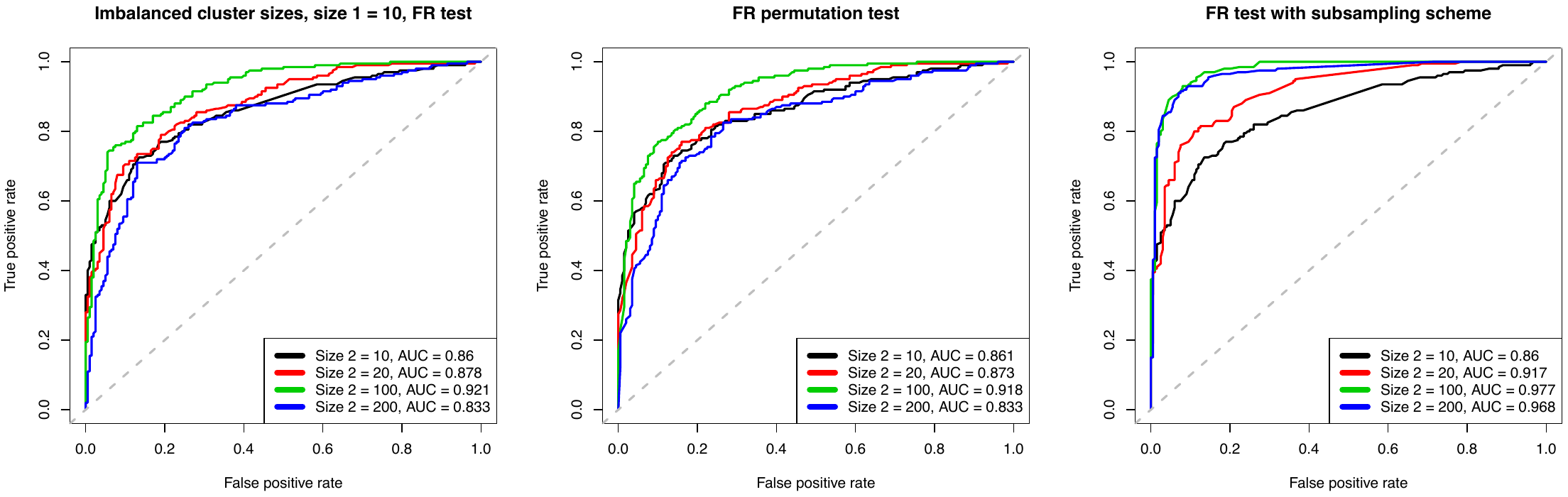

**Figure S30. Simulation for imbalanced cluster sizes.** ROC curves and AUC values for multivariate normal simulated data with size 1 fixed at 10 and size 2 varying from 10-200. Performance of the original FR test (left), FR permutation test (middle), and FR test with subsampling scheme (right) are compared with respect to the balanced/imbalance cluster sizes. The blue curve (size 1 = 10, size 2 = 200) show that FR test with the subsampling scheme (subsampling size = 10) outperforms the orginal FR and FR permutation tests.

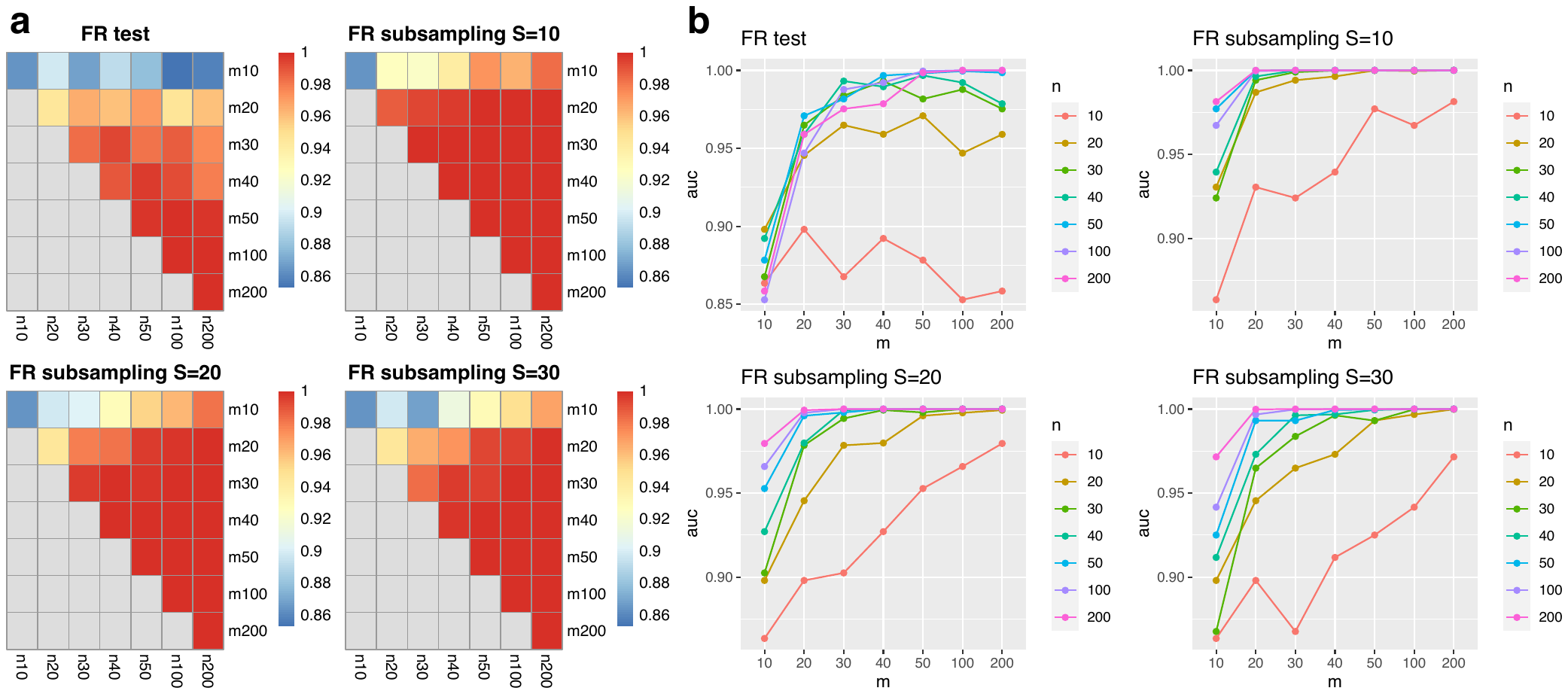

**Figure S31. Simulation performance of FR test and FR subsampling test with 𝑆 = 10, 20, 30. (a)** Heatmaps and **(b)** line plots show AUC values for the evaluated tests in simulated cases with cluster sizes varying from 20 to 200.

**Suppelementary Figure 32. Seurat results with cluster-level matching threshold at 40%.** Two-way matching results are shown in three colors: red indicates that a pair of clusters are matched in both directions (Layer 1 query to MTG reference with MTG markers, and MTG query to Layer 1 reference with Layer 1 markers); yellow indicates that a pair of clusters are matched in either direction; and blue indicates that a pair of clusters are not matched.

**Supplementary Table 1. Ensemble matching results across methods.** Reported are two-way matches agreed upon by two out of the three cell-to-cell, cell-to-cluster, and cluster-to-cluster matching methods.
